# Effort Drives Saccade Selection

**DOI:** 10.1101/2024.02.06.579052

**Authors:** Damian Koevoet, Laura Van Zantwijk, Marnix Naber, Sebastiaan Mathôt, Stefan Van der Stigchel, Christoph Strauch

## Abstract

What determines where to move the eyes? We recently showed that pupil size, a well-established marker of effort, also reflects the effort associated with making a saccade (’saccade costs’). Here we demonstrate saccade costs to critically drive saccade selection: when choosing between any two saccade directions, the least costly direction was consistently preferred. Strikingly, this principle even held during search in natural scenes in two additional experiments. When increasing cognitive demand experimentally through an auditory counting task, participants made fewer saccades and especially cut costly directions. This suggests that the eye-movement system and other cognitive operations consume similar resources that are flexibly allocated among each other as cognitive demand changes. Together, we argue that eye-movement behavior is tuned to adaptively minimize saccade-inherent effort.

## 1 Main

Humans make fast, ballistic eye movements, called saccades, to explore the rich visual world [1]. Saccades are executed approximately three to four times per second [2, 3]. Where to saccade is therefore one of the most frequent decisions the brain is faced with [4].

It is well established that the physical properties of the environment (bottom-up information) [5–9], the goals of the observer (top-down information) [10–13], and prior knowledge about a scene (selection history) [14, 15] drive where the eyes are moved. However, even when these factors are kept constant, there are many systematic biases in eye-movement behavior, such as a bias for cardinal compared with oblique saccade directions [16–21]. The presence of these biases suggests that additional factors must contribute to the decision of where to saccade, here referred to as ’saccade selection’. Recent evidence suggests that the effort involved with planning and executing (eye) movements may be one crucial factor in driving action selection [22–29]. Effort is thought to be minimized whenever possible [30, 31], likely because it is costly to spend inherently limited cognitive resources [22, 32]. We here use the term ’saccade cost’ to describe the intrinsic effort associated with planning and executing saccades. Although saccades are relatively affordable [i.e. not very costly; 1, 33], they are executed very often [2, 3] and therefore even small costs should add up over time [22, 34]. We here hypothesized that affordable saccades are preferred over costly saccades. This would be in line with recent evidence from computational models suggesting that saccade costs predict saccade behavior [e.g. 23, 25, 35, 36]. These studies either assumed saccade costs or indirectly inferred them from gaze behavior itself. However, no study has been able to quantify saccade costs (neuro-)physiologically, and therefore been able to directly test this hypothesis until recently.

We recently demonstrated that the effort of saccade planning can be measured with pupil size, which allows for a physiological quantification of saccade costs as long as low-level visual factors are controlled for [34]. Pupil size is an established marker of effort [37–45]. For instance, loading more in working memory or tracking more objects results in stronger pupil dilation [45–53]. Pupil size not only reflects cognitive (or mental) effort but also the effort of planning and executing movements [38, 54, 55]. We leveraged this to demonstrate that saccade costs can be captured with pupil size, and are higher for oblique compared with cardinal directions [34]. Here, we addressed whether saccade costs predict where to saccade.

We hypothesized that participants would prefer affordable over costly saccades to minimize effort expenditure. To test this, we first mapped out saccade costs across directions by measuring pupil size during saccade planning. To assess saccade preferences across the same directions, a subsequent free choice saccade task was employed. Previewing our results, saccade costs indeed predicted saccade preferences, as affordable directions were preferred over costly alternatives. Strikingly, this general principle even held when participants searched for targets in natural scenes in two additional experiments: saccade cost remained a fundamental driver of saccade selection. If saccades and other cognitive operations consume the same resources, this should reflect in adaptively changing saccade preferences in light of altering cognitive demands. We tested this idea experimentally by comparing saccade preferences with and without an auditory dual-task. As hypothesized, participants made fewer saccades overall under increased cognitive demand and especially cut the most costly directions. This provides convergent evidence that saccades are costly and rely at least in part on the same cognitive resources as other cognitively demanding operations.

## 2 Results

### 2.1 Saccade costs differ around the visual field

Twenty human participants planned and executed saccades in 36 different directions at a fixed amplitude (10°, Figure 1a, b). Pupil size was measured to index effort and thereby saccade cost during saccade planning (-150 ms until 170 ms around cue offset; also see Supplementary Figure 1). Replicating our previous findings, we found that pupil size differed across directions [34] (Figure 1c-f). We observed a larger pupil size during planning of oblique saccades compared with cardinal saccades (*β* = 7.662, *SE* = 1.957, *t* = 3.916, *p* < .001). Downward saccades were associated with a larger pupil size than upward saccades (*β* = .556, *SE* = .171, *t* = 3.261, *p* = .001), and we found a slightly larger pupil size for leftward compared with rightward saccades (*β* = .226, *SE* = .095, *t* = 2.388, *p* = .017). These effects were not mediated by differences in saccade properties, such as duration, amplitude, peak velocity, and landing precision (Figure 1e, f). Together, this shows that saccade costs differ as a function of direction, indicating that certain saccades are more costly than others.

**Figure 1:**
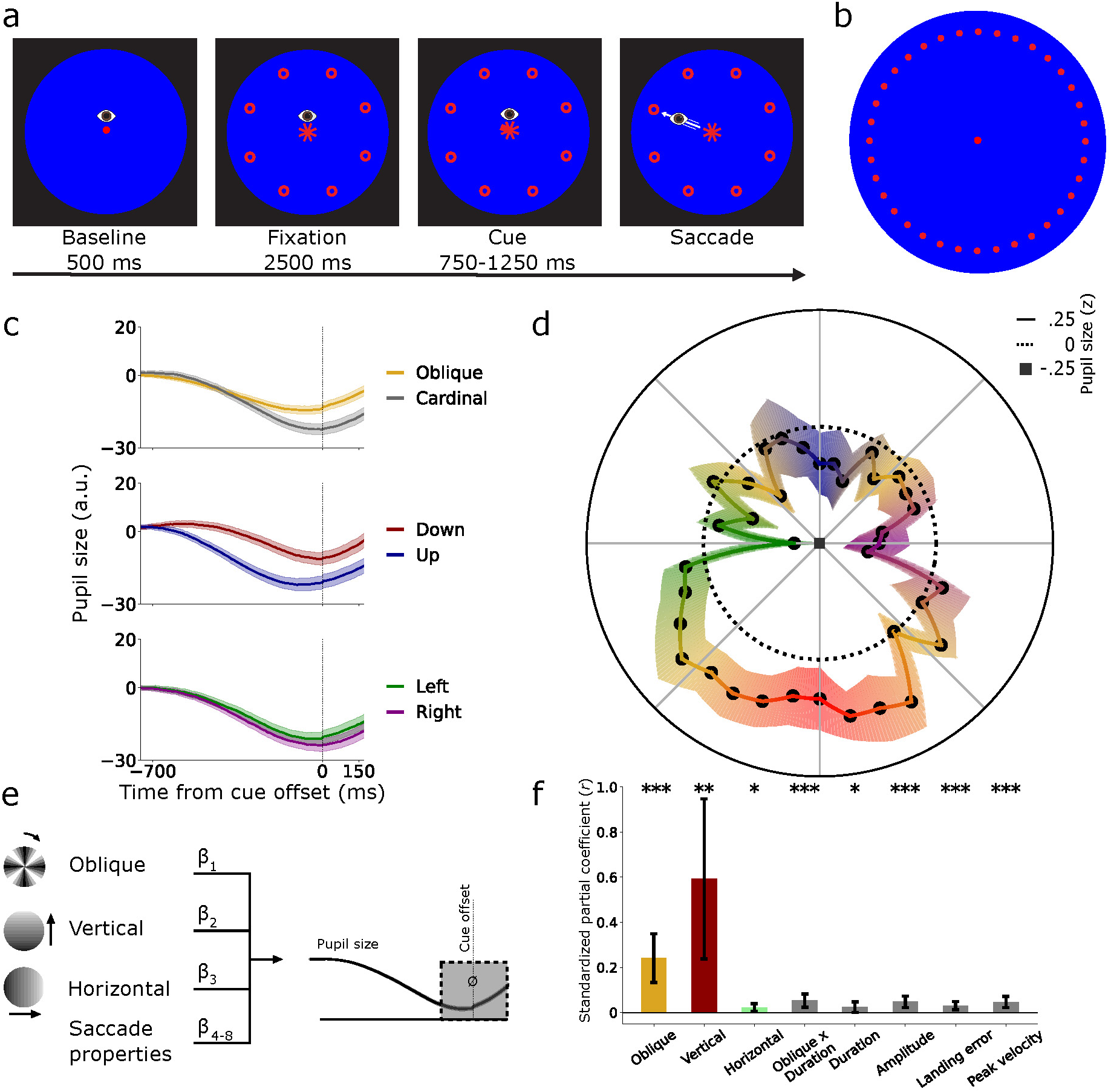
Pupil size differs during saccade planning across directions. **a** Twenty participants planned saccades in a cued direction. Saccades were executed as fast as possible upon cue offset. **b** All 36 possible saccade targets around the visual field. Only eight equally spaced locations were shown per trial. **c** Pupil size over time, split and averaged during saccade planning in oblique/cardinal, upward/downward, and left/rightward directions, locked to cue offset. Shaded areas indicate ± 1 s.e.m. **d** Averaged z-transformed pupil size during planning (-150 ms– 170 ms post cue, gray area in e) across directions. **e** Linear mixed-effects model using obliqueness, verticalness, horizontalness of directions, and saccade properties to predict pupil size during saccade planning. **f** Standardized partial coefficients per predictor with 95% confidence intervals. \**p* < .05, \*\**p* < .01, \*\*\**p* < .001.

### 2.2 Saccade costs predict saccade preferences

The same twenty participants subsequently completed a saccade preference task adapted from [23]. To determine which of the 36 saccade directions were preferred, participants freely chose between two possible saccade targets every trial (Figure 2a). We first analyzed whether saccade preferences differed across directions. Results showed that participants preferred cardinal over oblique directions (*β* = .091, *SE* = .023, *t* = 3.910, *p* < .001; Figure 2b), and preferred upward over downward directions (*β* = .036, *SE* = .017, *t* = 2.130, *p* = .033). No differences were observed between leftward and rightward saccade directions (*β* = .009, *SE* = .014, *t* = .668, *p* = .504).

These results indicate that saccade preferences seem to mirror the pattern of saccade costs (compare Figure 1d and Figure 2b). We proceeded to directly test if saccade costs predicted saccade preferences. To this end, we calculated the overall proportion that each direction was selected relative to how often it was offered to index saccade preferences (Figure 2b). In line with our hypothesis, pupil size during saccade planning (in cued directions) negatively correlated with saccade preferences (during self-selection) (*r*(35) = -.76, *p* < .001; Figure 2c). For the first time, this demonstrates that intrinsic saccade costs critically predict saccade preferences.

**Figure 2:**
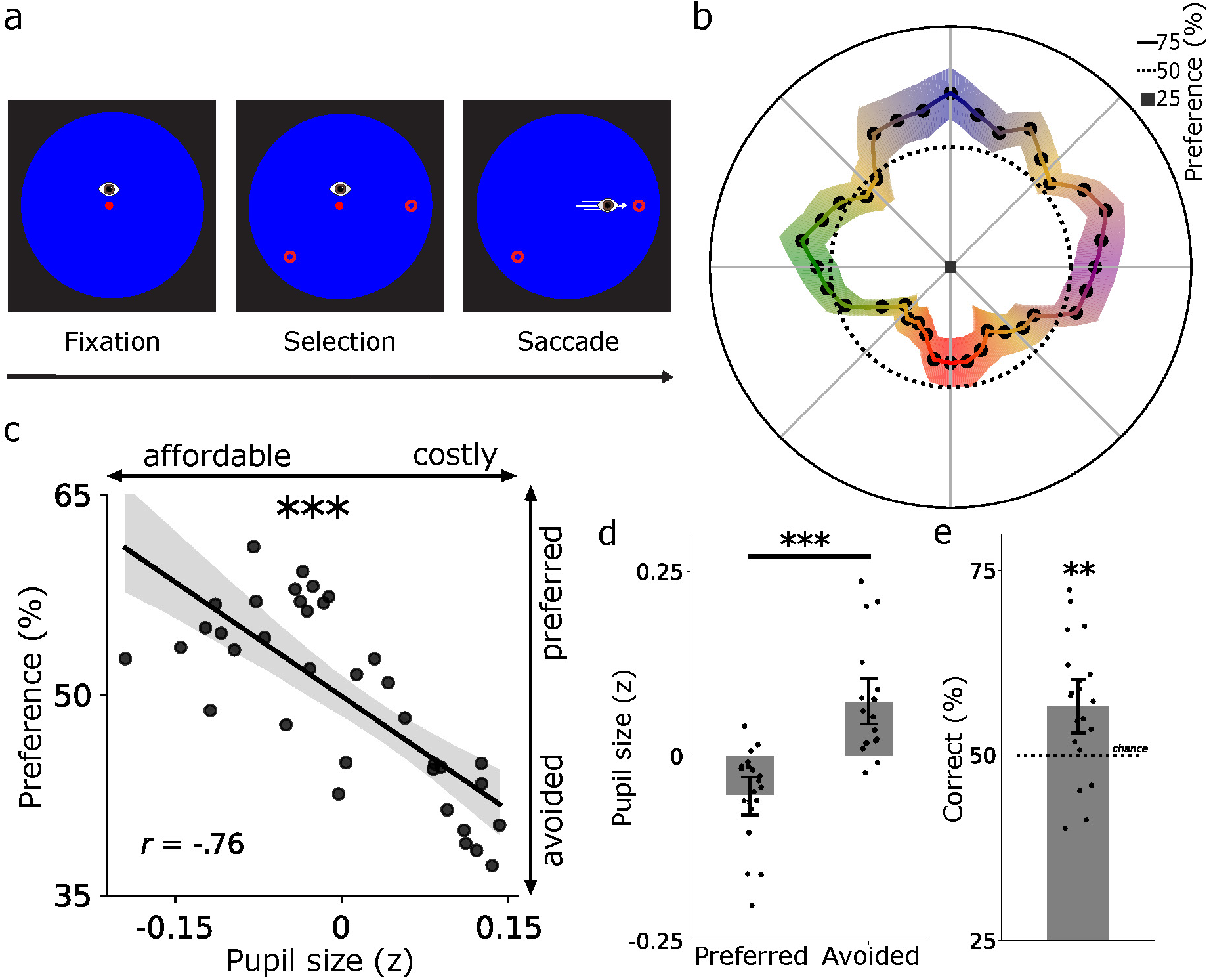
Saccade preferences differ across directions and are predicted by saccade costs. **a** The same twenty participants freely selected one of two saccade targets. **b** The average saccade preferences across directions (sum selected/sum offered). Shaded bands indicate ± 1 s.e.m.. **c** Saccade costs correlated negatively with saccade preferences across directions: costly directions were avoided and affordable directions preferred. Black datapoints represent directions (averaged across participants). **d** Pupil size was larger for avoided compared with preferred directions. **e** Saccade costs predicted saccade selection on a trial-by-trial basis (56.64%). Together, the saccade costs in the first task predicted saccade preferences in the subsequent task. **c-e** Error bars reflect bootstrapped 95% confidence intervals. **d-e** Black datapoints represent participants. \*\**p* < .01, \*\*\**p* < .001.

Put differently, participants preferred saccading towards affordable over costly options. A control analysis ruled out that the correlation between pupil size and saccade preferences was driven by other oculomotor metrics such as saccade latency and landing precision (see Supporting Information). Illustrating the robustness of this relationship further, we found smaller pupil sizes during saccade planning for preferred (selected >50%) than for avoided (selected <50%) directions (*t*(19) = 4.38, *p* < .001, Cohen’s *d* = .979; Figure 2d). As another test of the robustness of the effect, we analyzed whether saccade costs predicted saccade selection on a trial-by-trial basis. To this end, we first determined the more affordable option for each trial using the established saccade cost map (Figure 1d). We predicted that participants would select the more affordable option. Complementing the above analyses, the more affordable option was chosen above chance level across participants (*M* = 56.64%, 95%-CI = [52.75%-60.52%], one-sample *t*-test against 50%: *t*(19) = 3.26, *p* = .004, Cohen’s *d* = .729; Figure 2e). Together, these analyses established that saccade costs robustly predict saccade preferences.

### 2.3 Saccade costs predict saccade curvature and latency

If saccade cost is indeed weighed during saccade selection, this should be reflected in the oculomotor properties of the ensuing saccade. Saccade curvature reflects conflict between target and distractor saccade vectors: if a distractor is inhibited, and the target is activated, the saccade curves away from the distractor location [56–58]. Whenever saccade costs differ more between directions, there should therefore be more conflict between saccade vectors. If both directions were equally costly, there would be no need for conflict as cost minimization is impossible. We therefore hypothesized signs of increased oculomotor conflict to especially show in trials with relatively large differences in saccade costs. Furthermore, weighing costs (and reward) in decision-making is known to take time [59, 60]. More elaborate decisions should therefore not only show in more curvature, but also in longer saccade latencies.

**Figure 3:**
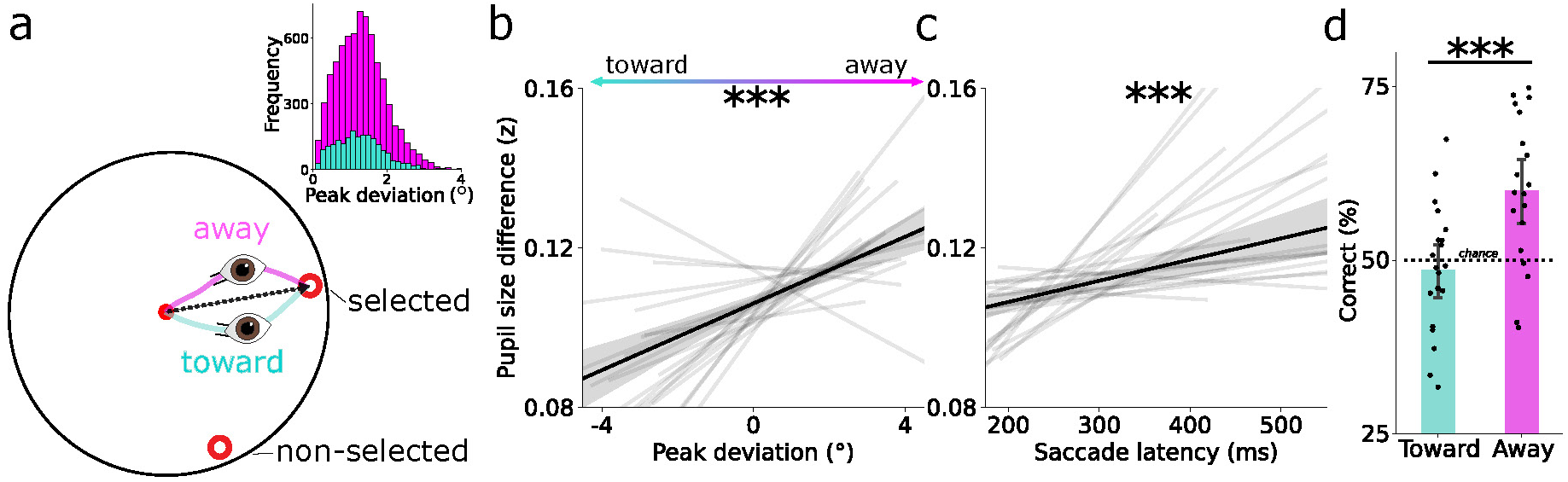
Saccade curvature and latency reveal active weighing of cost during saccade selection. **a** Schematic layout of saccade trajectories curving away (magenta) or toward (cyan) non-selected options. Curvature was calculated as the peak deviation from a straight line between gaze positions at saccade onset and offset. The top-right histogram shows that more saccades curved away than toward the non-selected option. **b** Difference in pupil size during the saccade planning task is linked to the peak curvature deviation in the saccade preference task. **c** Same as b, but now linked to saccade latency in the saccade preference task. Larger differences in pupil size are related to more oculomotor conflict between the two options, as reflected in more curvature away from the non-selected option and slower saccade latencies. **b, c** Black line depicts the relationship across all trials, gray lines denote regression fits per participant. **d** Saccade-cost based prediction of saccade selection split for toward and away curving trials. On a trial-by-trial basis, saccade costs predicted saccade selection above chance (59.72%) when saccades curved away from the non-selected option. In contrast, saccade costs did not predict saccade selection for ’toward’ saccades. Black datapoints represent participants. All error bars reflect bootstrapped 95% confidence intervals. \*\*\**p* < .001

To test this, we first split trials into saccades curving toward and away from the non-selected option. Saccades curved away from the non-selected option in the majority of trials, indicating oculomotor conflict (Figure 3a; *M* = 78.15%, 95%-CI = [74.734%-81.567%], *t*(19) = 15.741, *p* < .001, Cohen’s *d* = 3.519). We then examined how saccade curvature and latency predicted the difference in pupil size between the two possible saccade targets. Whenever the difference in pupil size between the two options was larger, saccades curved away more from the non-selected option (*β* = .004, *SE* = .001, *t* = 4.448, *p* < .001; Figure 3b), and their latencies slowed (*β* = .050, *SE* = .013, *t* = 4.323, *p* < .001; Figure 3c). Our results show that cost is actively weighed and leads to stronger conflict whenever cost differences are larger. This suggests that more elaborate decisions in saccade selection are predominantly made when warranted by sufficient differences in saccade costs between options.

The above analyses show that saccade costs affect oculomotor conflict, but does increased conflict between saccade vectors also lead to selecting more affordable options? We expected that especially when oculomotor conflict was high, participants would choose the more affordable option. This means that saccade costs should be more predictive of saccade selection in away compared with toward curving trials. To test this idea, we repeated the trial-by-trial prediction of which option was selected as before (Figure 2e), but now separately for trials with toward and away curving saccades. Pupil size (i.e. saccade cost) predicted saccade selection when saccades curved away (*M* = 59.72%, 95%-CI = [55.208%-64.233%], *t*(19) = 4.116, *p* < .001, Cohen’s *d* = .920), but not toward (*M* = 48.42%, 95%-CI = [44.496%-52.338%], *t*(19) = .771, *p* = .450, Cohen’s *d* = .172) the non-selected option. These prediction accuracies differed between curve directions (*t*(19) = 4.795, *p* < .001, Cohen’s *d* = 1.072; Figure 3d). This shows that saccade costs were predominantly considered when saccades curved away. Together, these analyses suggest that the costs of potential saccade targets are especially weighed during saccade selection when warranted by large differences in saccade costs. In these cases, oculomotor conflict increases and saccade cost plays a bigger role in saccade selection.

### 2.4 Saccade costs predict saccade preferences in natural viewing

The previous results establish that saccade costs predict saccade preferences in highly controlled settings. However, a crucial question is whether saccade costs also predict saccade preferences in more complex and less controlled settings, in which physical saliency, the observer’s goals, and prior knowledge about the scene also affect saccade selection. To test this, we analyzed data from two existing datasets [61] wherein participants (total *n* = 41) searched for small targets (‘Z’ or ‘H’) in natural scenes (Figure 4a; [62]). Again, we tested whether pupil size prior to saccades negatively linked with saccade preferences across directions. Because saccade costs and preferences across directions could differ for different situations (i.e. natural viewing vs. saccade preference task), but should always be negatively linked, we established both cost and preferences independently in each dataset. Many factors influence pupil size in such a natural task, for which we controlled as much as possible by including variables known to covary with pupil size in a linear mixed-effects model (based on [61]; e.g. luminance, gaze position, saccade properties, saliency, fixation number; see Methods) to access the underlying saccade costs. As hypothesized, we observed a negative relationship between pupil size and saccade preferences in both experiments (Exp. 1: *β* = 1.784, *SE* = .324, *t* = 5.412, *p* < .001; saccade preferences in Figure 4b, link in Figure 4c; Exp. 2: *β* = .644, *SE* = .170, *t* = 3.780, *p* < .001; saccade preferences in Figure 4d, link in Figure 4e). This shows that even when participants made unconstrained eye movements in natural scenes, saccade cost remained linked to saccade preferences: affordable directions were preferred over costly directions.

Do cognitive operations and eye movements consume from a similar pool of resources [45]? If so, increasing cognitive demand for non-oculomotor processes should result in decreasing available resources for the oculomotor system. In line with this idea, previous work indeed shows altered eye-movement behavior under effort as induced by dual tasks, for example by making less saccades under increased cognitive demand [63–65]. We therefore investigated whether less saccades were made as soon as participants had to count the occurrence of a specific digit in the auditory number stream in comparison to ignoring the stream (in Exp. 2; Figure 4a). Participants were instructed to prioritize the auditory digit-counting task over finding the visual search target. Therefore, resources should be shifted from the oculomotor system to the primary auditory counting task. The additional cognitive demand of the dual task indeed led to a decreased saccade frequency (*t*(24) = 7.224, *p* < .001, Cohen’s *d* = 1.445; Figure 4h). This indicates that the auditory dual task and the oculomotor system, at least in part, consumed from a shared pool of cognitive resources.

From a costs-perspective, it should be efficient to not only adjust the number of saccades (non-specific), but to also cut especially expensive directions the most (specific). Therefore, we expected participants to especially avoid costly saccades (as assessed in the single task) under higher cognitive demand (induced by the dual task). We calculated a saccade-adjustment map (Figure 4g) by subtracting the saccade preference map in the single task (Figure 4f) from the dual task map (Figure 4d). Participants seemingly cut vertical saccades in particular, and made more saccades to the top right direction. This pattern may have emerged as vertical saccades are more costly than horizontal saccades (also see Figure 1d). Oblique saccades may not have been cut because there were very little oblique saccades in the single condition to begin with (Figure 4d), making it difficult to observe a further reduction of such saccades under additional cognitive demand (i.e. a floor effect). Nevertheless, pupil size negatively linked with the adjustment map as hypothesized (*β* = 9.333, *SE* = .966, *t* = 9.659, *p* < .001; Figure 4i; while controlling for the same possible covariates as before). This shows that costly saccades were cut disproportionally when more cognitive resources were consumed by the additional auditory dual task. This demonstrates that cognitive resources are flexibly (dis)allocated from and to the oculomotor system based on the current resource demands.

## 3 Discussion

We here investigated whether effort determines saccade preferences. We first measured pupil size prior to saccade execution across directions as a physiological marker of effort and thus saccade costs. Next, saccade preferences were assessed in the same participants and directions. We observed that affordable saccades were preferred over costly ones. This is especially remarkable given that the delayed saccades in the planning task likely differ in their oculomotor program from the immediate saccades in the preference task in some regard. Furthermore, when two possible saccade directions differed more in saccade cost, we found higher oculomotor conflict as indexed by stronger saccade trajectory deviations away from the non-selected option and increased onset latencies. In two additional experiments, we demonstrated the link between saccade costs and saccade preferences to be robust even when participants made unconstrained eye movements during natural viewing. Lastly, saccade directions were flexibly adjusted based on cost as cognitive demand increased. Together, this demonstrates that saccade costs fundamentally underlie saccade selection, even when physical salience, the goals of the observer, and selection history affect where the eyes are moved.

**Figure 4:**
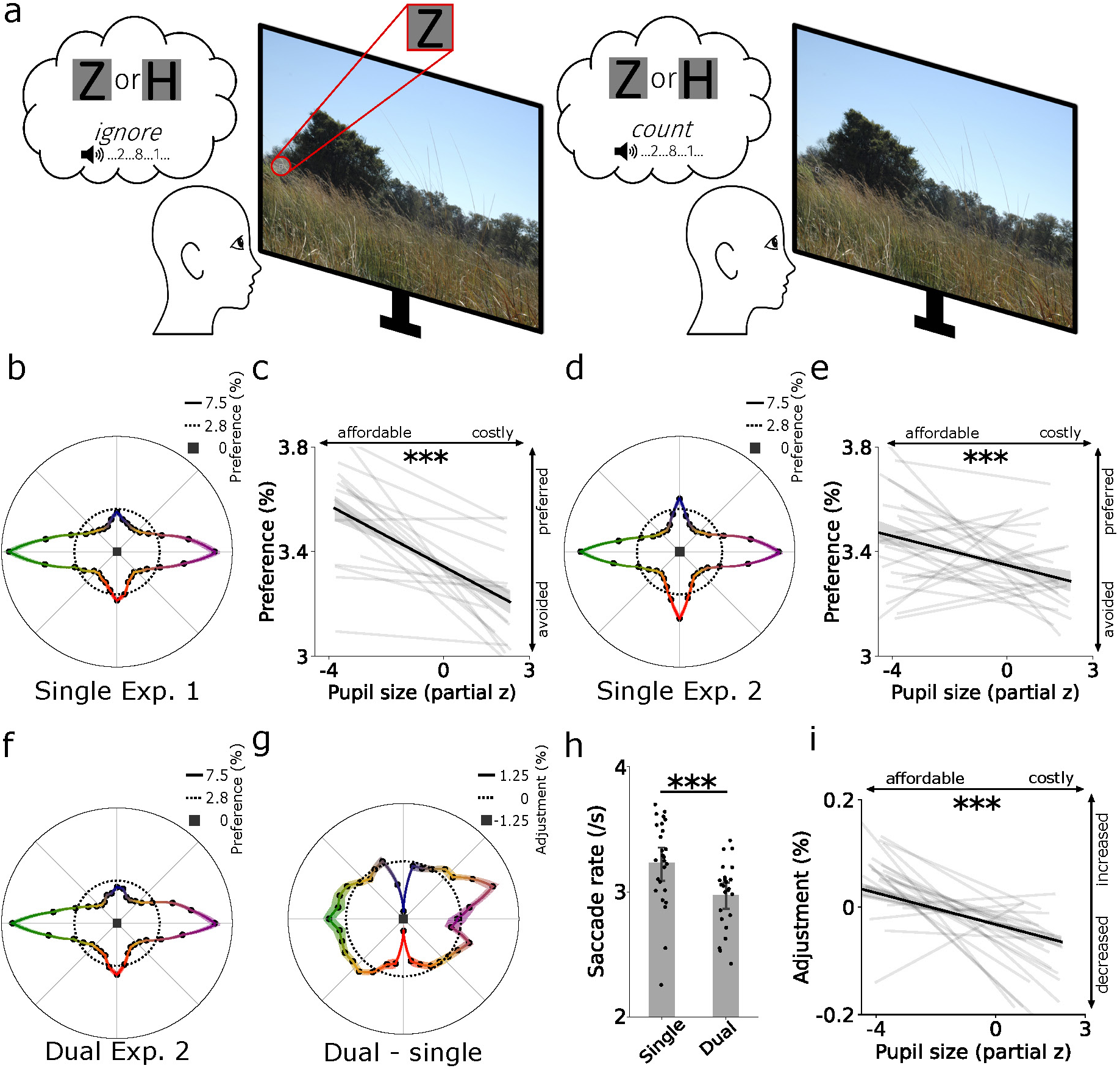
Saccade costs underlie saccade preferences in natural viewing. **a** Fourty-one participants searched for small letters (’Z’ or ‘H’) in natural scenes (Exp. 1; *n* = 16), and either ignored (single task) or additionally attended (dual task) to an auditory number stream (Exp. 2; *n* = 25). **b** Saccade preferences during search without auditory stimulation. **c** Preferred directions were associated with a smaller pupil size prior to the saccade (Exp. 1). **d, e** Same as b, c but now for Exp. 2 without attending the auditory number stream (single task). Preferred directions were again associated with a smaller pupil size preceding the saccade (Exp. 2). **f** Same as d but now under the increased cognitive demand of the (primary) auditory digit counting (dual) task. **g** Adjustment in saccade preferences between single- and dual-task conditions in percentage points. **h** Less saccades were executed in the more demanding dual-task condition. Black datapoints represent participants. **i** Pupil size during the single task predicted direction adjustments under additional cognitive demand. Costly saccades as assessed during the single-task condition were especially cut in the dual-task condition. **b, d, f, g** Shaded bands represent ±1 s.e.m.. Other error bars reflect bootstrapped 95% confidence intervals. **c, e, i** Black lines depict the relationship across all trials, gray lines denote regression fits per participant. \*\*\**p* < .001.

What contributes to intrinsic saccade costs? We speculate that at least three processes contribute to the total cost of a saccade [34]: the complexity of oculomotor programming [22, 66, 67], shifting of presaccadic attention [68, 69], and predictive/spatial remapping [70–73]. The complexity of an oculomotor program is arguably shaped by its neural underpinnings. For example, oblique but not cardinal saccades require communication between pontine and midbrain circuits [74–76]. Such differences in neural complexity may underlie the additional costs of oblique compared with cardinal saccades. Besides saccade direction, other properties of the ensuing saccade such as its speed, distance, curvature, and accuracy may contribute to a saccade’s total cost [22, 34, 54, 77, 78] but this remains to be investigated directly. Furthermore, presaccadic attention is shifted prior to each saccade to prepare the brain for the abrupt changes in retinal input resulting from saccades through spatial/predictive remapping [68–72, 79, 80]. This preparation for upcoming changes in retinal input consumes neurocognitive resources and therefore likely also contributes to saccade costs [22, 34]. To better understand saccade selection more generally, future work should elucidate which processes contribute to saccade costs, and how costs shape different (aspects of) saccades.

The observed differences in saccade costs across directions could be linked to established anisotropies in perception [81–87], attention [88–93], saccade characteristics [88, 89, 93, 94], and (early) visual cortex [95–99] [also see 100]. For example, downward saccades are more costly than upward saccades, which mimics a similar asymmetry in early visual areas wherein the upper visual field is relatively under-represented [95–99]; similarly stronger presaccadic benefits are found for down-compared with upward saccades [88, 89]. Moreover, upward saccades are more precise than downward saccades [94]. Future work should elucidate where saccade cost or the aforementioned anisotropies originate from and how they are related - something that pupil size alone cannot address.

We here measured cost as the degree of effort-linked pupil dilation. In addition to pupil size, other markers may also indicate saccade costs. For example, saccade latency has been proposed to index oculomotor effort [101], whereby saccades with longer latencies are associated with more oculomotor effort. This makes saccade latency a possible complementary marker of saccade costs (also see Supplementary Materials). Although relatively sluggish, pupil size is a valuable measure of attentional costs for (at least) two reasons. First, pupil size is a highly established as marker of effort, and is sensitive to effort more broadly than only in the context of saccades [37–46, 49]. Pupil size therefore allows to capture not only the costs of saccades, but also of covert attentional shifts [34], or shifts with other effectors such as head or arm movements [55, 102]. Second, as we have demonstrated, pupil size can measure saccade costs even when searching in natural scenes (Figure 4). During natural viewing, it is difficult to disentangle fixation duration from saccade latencies, complicating the use of saccade latency as a measure of saccade cost. Together, pupil size, saccade latency, and potential other markers of saccade cost could fulfill complementary roles in studying the role of cost in saccade selection.

Our findings are in line with established effort-based models that assume costs to be weighed against rewards during decision-making [103–108]. In such studies, reward and cognitive/physical effort are often parametrically manipulated to assess how much effort participants are willing to exert to acquire a given (monetary) reward [e.g. 109, 110]. Whereas this line of work manipulated the extrinsic costs and/or rewards of decision options (e.g. perceptual consequences of saccades [111, 112] or consequences associated with decision options), we here focus on the intrinsic costs of the movement itself (in terms of cognitive and physical effort). Relatedly, the intrinsic costs of arm movements are also considered during decision-making: biomechanically affordable movements are generally preferred over more costly ones [26–28]. We here extend these findings in two important ways. First, until now, the intrinsic costs of saccades and other movements have been inferred from gaze behavior itself or by using computational modelling [23, 25–28, 35, 36, 113]. In contrast, we directly measured cost physiologically using pupil size. Secondly, we show that physiologically measured saccade costs predict where saccades are directed in a controlled binary preference task, and even during natural viewing. Our findings could unite state-of-the-art computational models [e.g. 23, 25, 35, 36, 114] with physiological data, to directly test the role of saccade costs and ultimately further our understanding of saccade selection.

Throughout this paper, we have used cost in the limited context of saccades. However, cost-based decision-making may be a more general property of the brain [32, 37, 115–117]. Every action, be it physical or cognitive, is associated with an intrinsic cost, and pupil size is likely a general marker of this [45]. Note, however, that pupil dilation does not always reflect cost, as the pupil dilates in response to many sensory and cognitive factors which should be controlled for, or at least considered, when interpreting pupillometric data [e.g., see 40, 41, 43, 118]. Effort-linked pupil dilations are thought to be, at least in part, driven by activity in the brainstem locus coeruleus (LC) [41, 119–121] [but other neurotransmitters also affect pupil size, e.g. 122, 123]. Activity in LC with its widespread connections throughout the brain [121, 124–128] is considered to be crucial for the communication within and between neural populations and modulates global neural gain [129–133]. Neural firing is costly [22, 134], and therefore LC activity and pupil size are (neuro)physiologically plausible markers of cost [41]. Tentative evidence even suggests that continued exertion of effort (accompanied by altered pupil dilation) is linked to the accumulation of glutamate in the lateral prefrontal cortex [135], which may be a metabolic marker of cost [also see 117, 135, 136].

Besides the costs of increased neural activity when exerting more effort, effort should be considered costly for a second reason: Cognitive resources are limited. Therefore, any unnecessary resource expenditure reduces cognitive and behavioral flexibility [22, 32, 37, 117]. As a result, the brain needs to distribute resources between cognitive operations and the oculomotor system. We found evidence for the idea that such resource distribution is adaptive to the general level of cognitive demand and available resources: Increasing cognitive demand through an additional primary auditory dual task led to a lower saccade frequency, and especially costly saccades were cut. In this case, it is important to consider that the auditory task was the *primary task*, which should cause participants to distribute resources from the oculomotor system to the counting task. In other situations, more resources could be distributed to the oculomotor system instead, for example to discover new sources of reward [22, 137]. Adaptive resource allocation from, and to the oculomotor system parsimoniously explains a number of empirical observations. For example, higher cognitive demand is accompanied by smooth pursuits deviating more from to-be tracked targets [138], reduced (micro)saccade frequencies [Figure 4; 64, 65, 139, 140], and slower peak saccade velocities [141–143]. Relatedly, more precise saccades are accompanied with worse performance in a crowding task [94]. Furthermore, it has been proposed that saccade costs are weighed against other cognitive operations such as using working memory [34, 144–147]. How would the resources between the oculomotor system and cognitive tasks (like the auditory counting task) be related? One possibility is that both consume from limited working memory resources [148, 149]. Saccades are thought to encode target objects in a mandatory fashion into (visual) working memory [80], and the counting task requires participants to keep track of the auditory stream and maintain count of the instructed digit in working memory. However, the exact nature of which resources overlap between tasks remain open for future investigation [also see 150]. Together, we propose that cognitive resources are flexibly (dis)allocated to and from the oculomotor system based on the current demands to establish an optimal balance between performance and cost minimization.

How does the brain keep track of saccade costs, and which areas use it during saccade selection? Although our data do not allow direct inferences about the precise neural circuitry underlying the computations of oculomotor selection, oculomotor control is generally thought to be steered by a network encompassing the frontal eye field [151, 152], the supplementary eye field [153–155], the anterior cingulate cortex [156–159], the superior colliculus [41, 74, 160–164], and the cerebellum [56, 165–167]. These areas are not just associated with oculomotor control, but are all also thought to be crucial for decision-making processes [164, 168–178]. It is plausible that the weighing of saccade costs during saccade selection is performed by this oculomotor-decision making network, but other areas such as orbitofrontal cortex may also play a role [179, 180].

We report a combination of correlational and causal findings. Despite the correlational nature of some of our results, they consistently support the hypothesis that saccade costs predicts saccade selection [which we predicted previously, 34]. Causal evidence was provided by the dual-task experiment as saccade frequencies - and especially costly saccades were reduced under additional cognitive demand. Only a cost account predicts 1) a link between pupil size and saccade preferences, 2) a cardinal saccade bias, 3) reduced saccade frequency under additional cognitive demand, and 4) disproportional cutting of especially those directions associated with more pupil dilation. Together, our findings converge upon the conclusion that effort drives saccade selection.

To conclude, we have demonstrated that saccade costs can be measured using pupil size and that these costs robustly predict saccade selection. We propose that saccade selection is driven by physical properties of the environment, the observer’s goals, selection history, and another fundamental factor: effort.

## 4 Methods

### 4.1 Saccade planning and saccade preference tasks

#### 4.1.1 Participants

Twenty-two participants with normal or corrected-to-normal vision took part in the saccade planning and preference tasks across two sessions. One participant was excluded due to only finishing a single session, and another dataset was discarded due to not following task instructions (<50% included trials in the saccade planning task). Twenty participants were included in the analyses for the saccade planning, and saccade preference tasks (age: *M* = 24.00, range: [19–31], 12 women, 8 men). The current sample size was comparable with previous work investigating saccade costs [23, 34]. The total number of trials was substantially larger in the current dataset than in Koevoet *et al.* [34] (14,400 vs. 4,800), albeit from a slightly smaller number of participants (*n* = 20 vs. *n* = 24). Participants provided written informed consent before taking part, and were awarded monetary compensation or course credits. The experimental procedure was approved by Utrecht University’s ethical review board of the Faculty of Social Sciences (22-0635).

#### 4.1.2 Apparatus and stimuli

Gaze position and pupil size were recorded at 1000 Hz with an Eyelink 1000 desktop mount (SR Research, Ontario, Canada) in a brightness- and sound-attenuated laboratory. A chin- and forehead-rest limited head movements. Stimuli were presented using PsychoPy [v.2022.2.5; 181] on an ASUS ROG PG278Q monitor (2560 x 1440, 100 Hz) positioned 67.5 cm away from eye position. The eye-tracker was calibrated (9 points) at the beginning of each session, during each break, and whenever necessary throughout the experiment (same procedure for both tasks, see below).

Potential saccade targets were eight equally spaced out red rings (1° diameter) positioned at an eccentricity of 10° visual angle. The central fixation stimulus was a red eight-legged asterisk of which each leg pointed towards one of the possible saccade targets (1°). These stimuli were presented on a blue circle (12° diameter; 11.64 cd/m^2^); the remaining part of the screen was black (0.73 cd/m^2^) to ensure equal brightness across all 36 possible target locations (Figure 1a, b).

To control for low-level visual effects on pupil size, the red color of all stimuli was made equiluminant to the blue background colour using a flicker fusion calibration [as in 34]. A blue background (HSV: 240.1.1) was presented continuously while a central red circle (5° diameter) continuously flickered at 25 Hz. Participants adjusted the luminance of the red color by moving the mouse across the horizontal plane of the screen until the flickering was the least noticeable, and then clicked the left mouse button to confirm. This procedure was performed thrice, and the average luminance of the red colour was used for the fixation and target stimuli throughout the task. Participants completed the flicker fusion calibration preceding each task.

#### 4.1.3 Procedure

The experiment started with the saccade planning task [34], wherein participants planned saccades into 36 different cued directions. Each trial started when the central stimulus was fixated for 500 ms. After a fixation period (2000 ms) eight equally spaced potential saccade targets were presented (randomized which eight out of the 36 between trials). Afterwards, one of the eight legs of the asterisk became slightly thicker, cueing a saccade target (750-1250 ms, 100% valid). Participants planned and withheld an eye movement until cue offset, and then executed the saccade as fast as possible. Trials ended upon fixating the target stimulus (within 3°) for 500 ms. Whenever participants saccaded too early or to an incorrect location, red feedback text was presented (“too early”, “wrong location”). Each session consisted of 360 trials, preceded by ten practice trials. Participants could initiate a break whenever they wanted.

Participants subsequently completed a saccade preference task [adapted from 23]. Upon briefly fixating the central stimulus (10-500 ms), possible saccade targets were presented in two out of the 36 positions around the visual field. The only restriction was that the two targets should at least differ 20° in angle to ensure targets were sufficiently spaced out - and to limit saccade averaging [182]. Trials ended when one of the two saccade targets was fixated for 50 ms. Participants completed 360 trials per session.

#### 4.1.4 Data processing and analyses

All data were analyzed using custom Python (v3.9.14) and R (v4.3.1) scripts. Analyses of pupillometric data followed recommendations by [41, 118]. Blinks were interpolated [118], data were downsampled to 100 Hz, and pupil data were subtractively baseline corrected with the mean of the first 250 ms after cue onset. Saccades were detected offline using an onset velocity threshold of 75°/s and an offset threshold of 1°/s. Trials with fast (<175 ms) or slow (>550 ms) onset latency [34], a very short (<10 ms) or long saccade (>110 ms) duration [183], an amplitude smaller than 5°, or saccades landing more than 2° from the target location, saccades toward the wrong target, and practice trials were discarded (13.67% in total). To map out saccade planning costs across directions, the average pupil size was calculated 150 ms before until 170 ms after cue offset (Figure 1f) - before any saccade onsets to prevent pupil foreshortening errors [184]. We analyzed saccade costs by incorporating continuous predictors for oblique (cardinal vs. oblique; 0-4), vertical (up vs. down; in y coordinates) and horizontal (left vs. right; in x coordinates) direction biases in a linear mixed-effects model (Figure 1e, f). We also incorporated properties of the ensuing saccade to control for their possible associations with pupil size. The final model was determined using AIC-based backward model selection (Wilkinson notation: pupil size ∼ oblique*saccade duration + vertical + horizontal + amplitude landing error + peak velocity + (1 + oblique + vertical|participant)). For all mixed-effects models, we included as many by-participant random slopes as possible for our main effects of interest while ensuring model convergence [61, 185]. Absolute effect sizes (i.e. *r*) and their corresponding 95% confidence intervals for the linear mixed-effects models were calculated using *t* and *df* values with the ‘effectsize’ package (v.0.8.8) in R. To obtain an average saccade costs map, pupil sizes were z-transformed per participant within sessions, and then averaged across participants for each direction (Figure 1d).

For the saccade preference task, saccades were detected as above. Which saccade target was selected per trial was determined using the last 50 ms of gaze data of each trial - the option closest to the gaze position was treated as the selected target. Trials were discarded if the difference in distance between the two saccade options in gaze position was less than 1.5° (3.47%). A logistic mixed-effects model was fit to investigate anisotropies across directions (Wilkinson notation: saccade preference ∼ oblique + vertical + horizontal + (1 + oblique + vertical + horizontal|participant)). Saccade preference for each direction was calculated per participant by summing how often a direction was chosen and then dividing by the number of times that direction was offered. As for the saccade cost map, the average saccade preference map was obtained by averaging across participants (Figure 2b). Preferred (>50%) and avoided (<50% chosen) directions were grouped using the average preference map (Figure 2d). To investigate if cost predicted saccade selection on a trial-by-trial basis, we compared the saccade costs of the two potential options. We predicted that the option with the smaller pupil size from the average cost map (obtained from the saccade planning task) would be chosen. This procedure was performed for each participant, and subsequently tested against chance performance (50%) with a one-sample *t*-test.

Saccade curvature was computed using the peak deviation from a straight line between gaze position at saccade onset until saccade offset [Figure 3a; 56]. Trials were excluded from these analyses if: a) saccade latencies were shorter than 175 ms, b) saccade amplitudes were smaller than 5°, c) saccade durations were shorter than 10 ms or longer than 110 ms and d) if the angle between targets exceeded 150° [34, 182, 183]. To analyze the relationship between saccade costs and saccade properties, we first computed the absolute saccade cost difference for each trial (as indexed from the average saccade cost map). A linear mixed-effects model was conducted to test whether saccade curvature and latency linked to saccade costs (Wilkinson notation: cost difference ∼ peak deviation + latency + (1 + peak deviation|participant)).

We then split the data based on whether saccades curved toward or away from the non-selected option. The same trial-by-trial analysis as described above was used to investigate if cost predicted saccade selection in toward and away trials separately.

### 4.2 Search in natural scenes

#### 4.2.1 Procedure

We analyzed existing data of two experiments to investigate if effort drives saccade selection in a more natural task [for an exhaustive explanation of the procedure see the original paper 61]. Briefly, in Experiments 1 and 2, sixteen and twenty-five participants respectively searched for small letters (‘Z’ or ‘H’) in natural scenes (from [62]). As in the saccade planning and preference tasks, gaze position and pupil size were recorded with an Eyelink 1000 (SR Research, Ontario, Canada) at 1000 Hz. Stimuli were presented (1280 x 1024) using OpenSesame [186] with the PsychoPy backend [181].

In Experiment 2, auditory digits (0-9) were presented with an inter-digit interval of 1500 ms during search - note that Experiment 1 did not feature the auditory dual task. Crucially, participants either performed a dual task wherein the count of a specific digit was monitored throughout search, or a single task where the number stream was ignored. The single and dual conditions were blocked, and the sequence of these blocks was random across participants.

#### 4.2.2 Data processing and analyses

Pupil size was averaged per fixation and subsequently z-transformed per participant [61]. Fixations with pupil sizes deviating more than 3*SD* from the mean (within a participant) and fixations positioned outside of the monitor were excluded to mitigate possible confounds (Exp. 1: 4.32%, Exp. 2: 9.75% discarded). 57,127 and 214,449 fixations were analyzed from Experiments 1 and 2, respectively. Fixations were classified into 36 bins based on their direction (bins consistent with the saccade planning and preference tasks).

To investigate if saccade costs predicted saccade preferences when searching in natural scenes, we analyzed all fixations from Experiment 1, and fixations from the single condition in Experiment 2. Next, we computed the average saccade preference map separately for each experiment by calculating the percentage of saccades in any of the 36 directions. Linear mixed-effects models were used to investigate whether this preference map predicted pupil size on a fixation-by-fixation basis in both experiments. We controlled for as many possible factors that are known to covary with pupil size in our model to control for them as much as possible to attempt to access the underlying saccade cost signal [61]; Wilkinson notation: pupil size ∼ saccade preferences + luminance + saliency + fixation number + trial number + x gaze coordinate + y gaze coordinate + saccade duration + fixation duration + saccade amplitude + (1 + saccade preferences|participant).

To investigate if costly saccades were avoided in particular when the overall level of demand increased via the dual task, we analyzed data from Experiment 2, The percentages of saccades made into each direction for the single and dual conditions were calculated. We subtracted these averaged preference maps to obtain an adjustment map: this revealed how participants altered their saccade preferences under additional demand (Figure 4e). We predicted pupil size using the average adjustment map for each direction while again controlling for many possible confounding factors in the single condition using a linear mixed-effects model on a fixation-by-fixation basis (Wilkinson notation: pupil size ∼ saccade adjustment + luminance + saliency + fixation number + trial number + x gaze coordinate + y gaze coordinate + saccade duration + fixation duration + saccade amplitude + (1 + saccade adjustment|participant)).

## Acknowledgments

This project has received funding from the European Research Council (ERC) under the European Union’s Horizon 2020 research and innovation programme (grant agreement n° 863732).

## Data availability and code availability

Data and analyses scripts to reproduce the results are available via the Open Science Framework: https://osf.io/n3ktm/.

## Supporting Information

**Supporting Figure 1:**
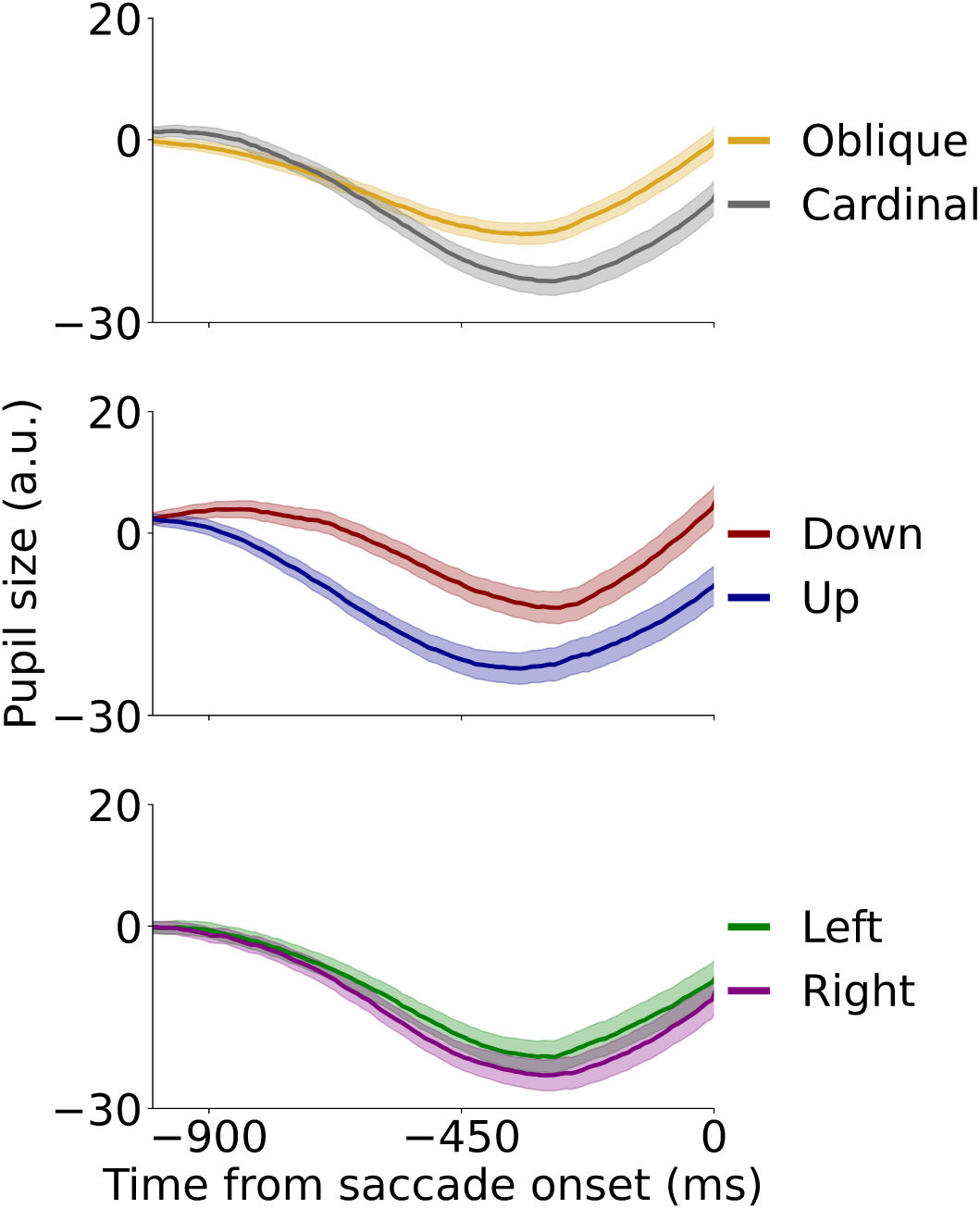
Saccade-locked pupil traces as index of saccade costs in different directions. Using the average pupil size in the 350 ms before saccade onset (saccade-locked), results remained qualitatively identical. Planning oblique saccades was associated with a larger pupil size than cardinal ones (*β* = 9.897, *SE* = 2.223, *t* = 4.451, *p* < .001), and downward saccades were associated with a larger pupil size than upward saccades (*β* = .471, *SE* = .112, *t* = 4.189, *p* < .001). A slightly increased pupil size for leftward compared with rightward saccades was observed as well (*β* = .260, *SE* = .107, *t* = 2.436, *p* = .015).

### Supporting Analysis

To ascertain whether pupil size or other oculomotor metrics predict saccade preferences, we conducted a multiple regression analysis. We calculated average pupil size, saccade latency, landing precision and peak velocity maps across all 36 directions. The model, determined using AIC-based backward selection, included pupil size, latency and landing precision as predictors (Wilkinson notation: saccade preferences ∼ pupil size + saccade latency + landing precision). The analysis revealed that pupil size (*β* = -42.853, *t* = 4.791, *p* < .001) and saccade latency (*β* = -.377, *t* = 2.106, *p* = .043) predicted saccade preferences. Landing precision did not reach significance (*β* = 23.631, *t* = 1.675, *p* = .104). Together, this demonstrates that although other oculomotor metrics such as saccade latency contribute to saccade selection, pupil size remains a robust marker of saccade selection.

**Supporting Table 1:** Full outcomes of the linear mixed-effects model analyzing pupil size assessed saccade costs across directions.

| Predictor | $\beta$ | SE | $t$ | $p$ |
| --- | --- | --- | --- | --- |
| Intercept | -26.510 | 9.059 | -2.926 | 0.003 |
| Obliqueness | 7.662 | 1.957 | 3.916 | < 0.001 |
| Verticalness | -0.556 | 0.171 | -3.261 | 0.001 |
| Horizontalness | -0.226 | 0.095 | -2.388 | 0.017 |
| Obliqueness x Duration | -0.109 | 0.031 | -3.495 | < 0.001 |
| Duration | 0.169 | 0.085 | 1.981 | 0.048 |
| Amplitude | -2.423 | 0.656 | -3.691 | < 0.001 |
| Landing error | 7.344 | 2.168 | 3.387 | 0.001 |
| Peak velocity | 0.054 | 0.015 | 3.692 | < 0.001 |
| Participant Var | 602.851 | 2.786 |  |  |
| Participant x Obliqueness Cov | -55.890 | 0.395 |  |  |
| Participant x Verticalness Cov | -1.625 | 0.058 |  |  |
| Obliqueness x Verticalness Cov | 0.934 | 0.011 |  |  |
| Obliqueness Var | 13.790 | 0.083 |  |  |
| Verticalness Var | 0.380 | 0.003 |  |  |

**Supporting Table 2:** Full outcomes of the linear mixed-effects model predicting pupil size using saccade preferences and control variables in Experiment 1.

| Predictor | $\beta$ | SE | *t* | *p* |
| --- | --- | --- | --- | --- |
| Intercept | 0.825 | 0.061 | 13.504 | < 0.001 |
| Direction preferences | -1.784 | 0.324 | -5.412 | < 0.001 |
| X coordinate | -0.0002 | 0.00001 | -21.12 | < 0.001 |
| Y coordinate | -0.0002 | 0.00001 | -11.316 | < 0.001 |
| Luminance | -0.635 | 0.017 | -36.251 | < 0.001 |
| Saliency | -0.001 | 0.0001 | -10.536 | < 0.001 |
| Trial number | -0.006 | 0.00006 | -91.139 | < 0.001 |
| Fixation number (in trial) | 0.006 | 0.0002 | 55.513 | < 0.001 |
| Saccade duration | -0.004 | 0.0009 | -42.150 | < 0.001 |
| Fixation duration | 0.0003 | 0.00004 | 7.105 | < 0.001 |
| Amplitude | 0.009 | 0.0007 | 13.979 | < 0.001 |
| Participant Var | 0.053 |  |  |  |
| Direction preferences Var | 0.833 | 0.646 |  |  |
| Participant x Direction preferences Cov | 0.003 | 0.250 |  |  |

**Supporting Table 3:** Full outcomes of the linear mixed-effects model predicting pupil size using saccade preferences and control variables in Experiment 2.

| Predictor | $\beta$ | SE | *t* | *p* |
| --- | --- | --- | --- | --- |
| Intercept | 1.093 | 0.060 | 18.301 | < 0.001 |
| Direction preferences | -0.644 | 0.170 | -3.780 | < 0.001 |
| X coordinate | -0.0001 | 0.000005 | -24.442 | < 0.001 |
| Y coordinate | -0.00004 | 0.000007 | -5.072 | < 0.001 |
| Luminance | -0.341 | 0.009 | -38.269 | < 0.001 |
| Saliency | -0.001 | 0.00006 | -10.405 | < 0.001 |
| Trial number | -0.007 | 0.00006 | -126.687 | < 0.001 |
| Fixation number (in trial) | 0.006 | 0.00009 | 64.174 | < 0.001 |
| Saccade duration | -0.007 | 0.00005 | -152.775 | < 0.001 |
| Fixation duration | 0.000005 | 0.00002 | 0.732 | 0.464 |
| Amplitude | 0.013 | 0.0003 | 37.737 | < 0.001 |
| Participant Var | 0.086 | 0.041 |  |  |
| Direction preferences Var | 0.322 | 0.339 |  |  |
| Participant x Direction preferences Cov | 0.034 | 0.086 |  |  |

**Supporting Table 4:**
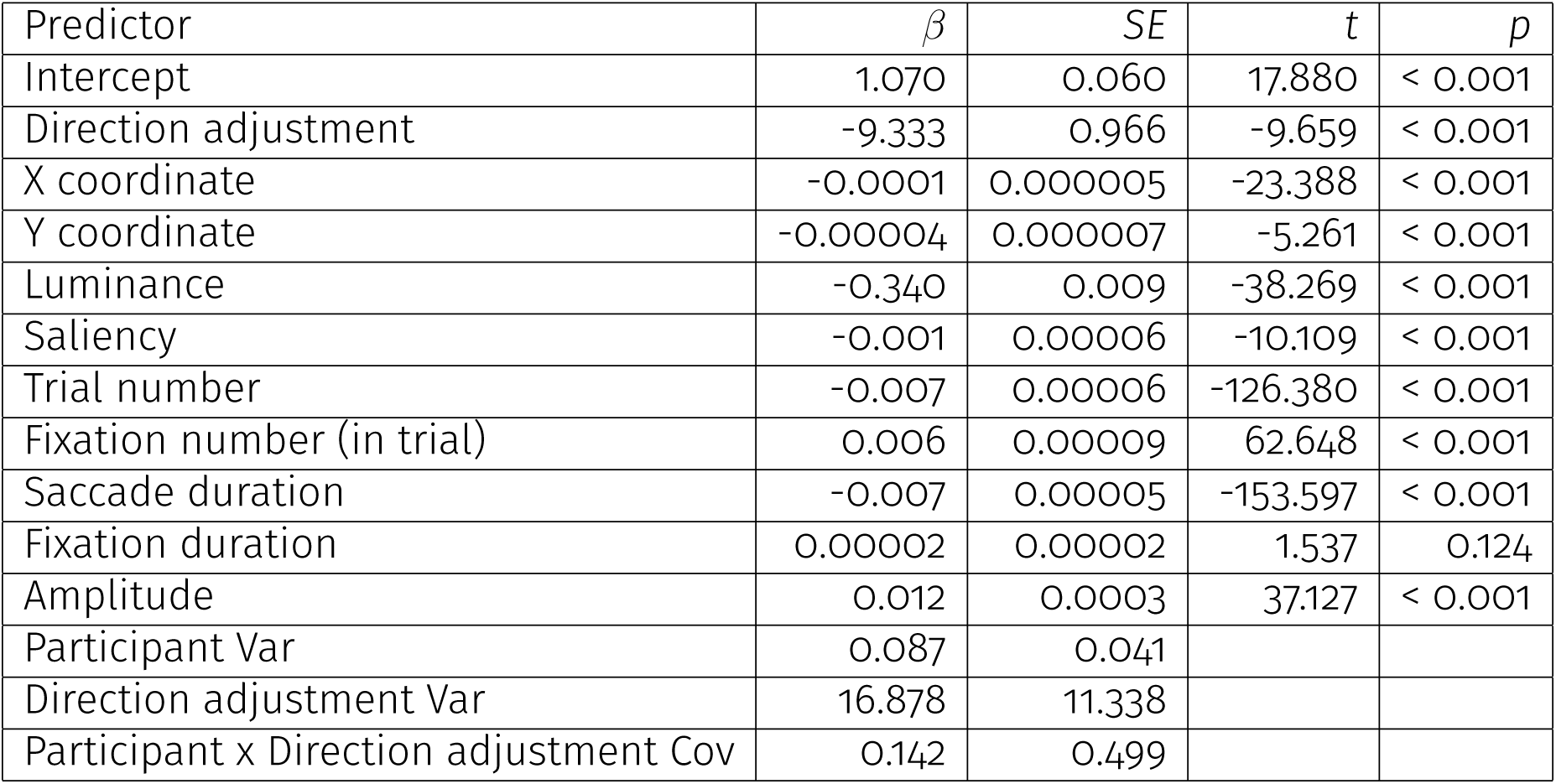
Full outcomes of the linear mixed-effects model predicting pupil size using saccade direction adjustment and control variables.

## References

1. 1. Findlay, J. M. & Gilchrist, I. D. Active Vision: The Psychology of Looking and Seeing isbn: 978-0-19-852479-3 (Oxford University Press, Aug. 2003).

2. Henderson, J. M. Human Gaze Control during Real-World Scene Perception. Trends in Cognitive Sciences 7, 498–504. issn: 1364-6613. (2022) (Nov. 2003).

3. Henderson, J. M. & Hollingworth, A. in Eye Guidance in Reading and Scene Perception (ed Underwood, G.) 269–293 (Elsevier Science Ltd, Amsterdam, Jan. 1998). isbn: 978-0-08-043361-5. (2023).

4. Bargary, G. et al. Individual Differences in Human Eye Movements: An Oculomotor Signature? Vision Research. Individual Differences as a Window into the Structure and Function of the Visual System 141, 157–169. issn: 0042-6989. (2023) (Dec. 2017).

5. Kümmerer, M., Bethge, M. & Wallis, T. S. A. DeepGaze III: Modeling Free-Viewing Human Scanpaths with Deep Learning. Journal of Vision 22, 7. issn: 1534-7362. (2023) (Apr. 2022).

6. Kümmerer, M., Wallis, T. S. A. & Bethge, M. DeepGaze II: Reading Fixations from Deep Features Trained on Object Recognition Oct. 2016. arXiv: 1610.01563 [cs, q-bio, stat]. (2023).

7. Itti, L., Koch, C. & Niebur, E. A Model of Saliency-Based Visual Attention for Rapid Scene Analysis. IEEE Transactions on Pattern Analysis and Machine Intelligence 20, 1254–1259. issn: 1939-3539. (2023) (Nov. 1998).

8. Itti, L. & Koch, C. Computational Modelling of Visual Attention. Nature Reviews Neuroscience 2, 194–203. issn: 1471-0048. (2023) (Mar. 2001).

9. Theeuwes, J. Stimulus-Driven Capture and Attentional Set: Selective Search for Color and Visual Abrupt Onsets. Journal of Experimental Psychology: Human Perception and Performance 20, 799–806. issn: 1939-1277 (1994).

10. Posner, M. I. Orienting of Attention. Quarterly Journal of Experimental Psychology 32, 3–25. issn: 0033-555X. (2022) (Feb. 1980).

11. Petersen, S. E. & Posner, M. I. The Attention System of the Human Brain: 20 Years After. Annual review of neuroscience 35, 73–89. issn: 0147-006X. (2022) (July 2012).

12. Posner, M. I. & Petersen, S. E. The Attention System of the Human Brain. Annual Review of Neuroscience 13, 25–42. issn: 1545-4126 (1990).

13. Desimone, R. & Duncan, J. Neural Mechanisms of Selective Visual Attention. Annual Review of Neuroscience 18, 193–222. issn: 1545-4126 (1995).

14. Awh, E., Belopolsky, A. V. & Theeuwes, J. Top-down versus Bottom-up Attentional Control: A Failed Theoretical Dichotomy. Trends in Cognitive Sciences 16, 437–443. issn: 1879-307X (Aug. 2012).

15. Theeuwes, J., Bogaerts, L. & van Moorselaar, D. What to Expect Where and When: How Statistical Learning Drives Visual Selection. Trends in Cognitive Sciences 26, 860–872. issn: 1364-6613. (2022) (Oct. 2022).

16. Foulsham, T. & Kingstone, A. Asymmetries in the Direction of Saccades during Perception of Scenes and Fractals: Effects of Image Type and Image Features. Vision Research 50, 779–795. issn: 0042-6989. (2022) (Apr. 2010).

17. Gilchrist, I. D. & Harvey, M. Evidence for a Systematic Component within Scan Paths in Visual Search. Visual Cognition 14, 704–715. issn: 1350-6285. (2022) (Aug. 2006).

18. Bays, P. M. & Husain, M. Active Inhibition and Memory Promote Exploration and Search of Natural Scenes. Journal of Vision 12, 8. issn: 1534-7362. (2023) (Aug. 2012).

19. Anderson, A. J., Yadav, H. & Carpenter, R. H. S. Directional Prediction by the Saccadic System. Current biology: CB 18, 614–618. issn: 0960-9822 (Apr. 2008).

20. Engbert, R. & Kliegl, R. Microsaccades Uncover the Orientation of Covert Attention. Vision Research 43, 1035–1045. issn: 0042-6989. (2022) (Apr. 2003).

21. Tatler, B. W. & Vincent, B. T. The Prominence of Behavioural Biases in Eye Guidance. Visual Cognition 17, 1029–1054. issn: 1350-6285. (2023) (Aug. 2009).

22. 22. Shadmehr, R. & Ahmed, A. A. Vigor: Neuroeconomics of Movement Control isbn: 978-0-262-04405-9 (MIT Press, July 2020).

23. Thomas, T., Hoppe, D. & Rothkopf, C. A. The Neuroeconomics of Individual Differences in Saccadic Decisions June 2022. (2023).

24. Hagura, N., Haggard, P. & Diedrichsen, J. Perceptual Decisions Are Biased by the Cost to Act. eLife 6 (ed Gold, J. I.) e18422. issn: 2050-084X. (2023) (Feb. 2017).

25. Kadner, F., Thomas, T., Hoppe, D. & Rothkopf, C. A. Improving Saliency Models’ Predictions of the next Fixation with Humans’ Intrinsic Cost of Gaze Shifts Sept. 2022. arXiv: 2207.04250 [cs]. (2023).

26. Cos, I., Duque, J. & Cisek, P. Rapid Prediction of Biomechanical Costs during Action Decisions. Journal of Neurophysiology 112, 1256–1266. issn: 0022-3077. (2024) (Sept. 2014).

27. Cos, I., Bélanger, N. & Cisek, P. The Influence of Predicted Arm Biomechanics on Decision Making. Journal of Neurophysiology 105, 3022–3033. issn: 0022-3077. (2024) (June 2011).

28. Cos, I., Medleg, F. & Cisek, P. The Modulatory Influence of End-Point Controllability on Decisions between Actions. Journal of Neurophysiology 108, 1764– 1780. issn: 0022-3077. (2024) (Sept. 2012).

29. Todorov, E. & Jordan, M. I. Optimal Feedback Control as a Theory of Motor Coordination. Nature Neuroscience 5, 1226–1235. issn: 1546-1726. (2024) (Nov. 2002).

30. Hull, C. L. Principles of Behavior: An Introduction to Behavior Theory x, 422 (Appleton-Century, Oxford, England, 1943).

31. Tsai, L. S. The Laws of Minimum Effort and Maximum Satisfaction in Animal Behavior. National Research Institute of Psychology (1932).

32. Friston, K. The Free-Energy Principle: A Unified Brain Theory? Nature Reviews Neuroscience 11, 127–138. issn: 1471-0048. (2022) (Feb. 2010).

33. Theeuwes, J. in The Influence of Attention, Learning, and Motivation on Visual Search (eds Dodd, M. D. & Flowers, J. H.) 23–62 (Springer, New York, NY, 2012). isbn: 978-1-4614-4794-8. (2022).

34. Koevoet, D., Strauch, C., Naber, M. & Van der Stigchel, S. The Costs of Paying Overt and Covert Attention Assessed With Pupillometry. Psychological Science 34, 887–898. issn: 0956-7976. (2023) (June 2023).

35. Hoppe, D. & Rothkopf, C. A. Learning Rational Temporal Eye Movement Strategies. Proceedings of the National Academy of Sciences 113, 8332–8337. (2024) (July 2016).

36. Hoppe, D. & Rothkopf, C. A. Multi-Step Planning of Eye Movements in Visual Search. Scientific Reports 9, 144. issn: 2045-2322. (2024) (Jan. 2019).

37. Kahneman, D. Attention and Effort isbn: 978-0-13-050518-7 (Prentice-Hall, 1973).

38. Bumke, O. Die Pupillenstörungen Bei Geistes-und Nervenkrankheiten 2nd ed. (Fischer, 1911).

39. Laeng, B., Sirois, S. & Gredebäck, G. Pupillometry: A Window to the Preconscious? Perspectives on Psychological Science 7, 18–27. issn: 1745-6916. (2022) (Jan. 2012).

40. Mathôt, S. Pupillometry: Psychology, Physiology, and Function. Journal of Cognition 1, 16. issn: 2514-4820. (2022) (2018).

41. Strauch, C., Wang, C.-A., Einhäuser, W., Van der Stigchel, S. & Naber, M. Pupillometry as an Integrated Readout of Distinct Attentional Networks. Trends in Neurosciences 45, 635–647. issn: 0166-2236. (2022) (June 2022).

42. Loewenfeld, I. E. The Pupil : Anatomy, Physiology, and Clinical Applications 1st ed. (2024) (Iowa State University Press, 1993).

43. Sirois, S. & Brisson, J. Pupillometry. WIREs Cognitive Science 5, 679–692. issn: 1939-5086 (2014).

44. van der Wel, P. & van Steenbergen, H. Pupil Dilation as an Index of Effort in Cognitive Control Tasks: A Review. Psychonomic Bulletin & Review 25, 2005– 2015. issn: 1531-5320. (2021) (Dec. 2018).

45. Beatty, J. Task-Evoked Pupillary Responses, Processing Load, and the Structure of Processing Resources. Psychological Bulletin 91, 276–292. issn: 1939-1455 (1982).

46. Koevoet, D., Strauch, C., Van der Stigchel, S., Mathôt, S. & Naber, M. Revealing Visual Working Memory Operations with Pupillometry: Encoding, Maintenance, and Prioritization. WIREs Cognitive Science, e1668. issn: 1939-5086. (2023) (2024).

47. Robison, M. K. & Unsworth, N. Pupillometry Tracks Fluctuations in Working Memory Performance. *Attention, Perception*, & Psychophysics 81, 407–419. issn: 1943-393X. (2021) (Feb. 2019).

48. Alnæs, D. et al. Pupil Size Signals Mental Effort Deployed during Multiple Object Tracking and Predicts Brain Activity in the Dorsal Attention Network and the Locus Coeruleus. Journal of Vision 14, 1. issn: 1534-7362. (2023) (Apr. 2014).

49. Kahneman, D. & Beatty, J. Pupil Diameter and Load on Memory. Science (New York, N.Y.) 154, 1583–1585. issn: 0036-8075 (Dec. 1966).

50. Unsworth, N. & Miller, A. L. Individual Differences in the Intensity and Consistency of Attention. Current Directions in Psychological Science 30, 391–400. issn: 0963-7214. (2023) (Oct. 2021).

51. Unsworth, N. & Robison, M. K. Individual Differences in the Allocation of Attention to Items in Working Memory: Evidence from Pupillometry. Psychonomic Bulletin & Review 22, 757–765. issn: 1531-5320. (2022) (June 2015).

52. Ahern, S. & Beatty, J. Pupillary Responses During Information Processing Vary with Scholastic Aptitude Test Scores. Science 205, 1289–1292. (2022) (Sept. 1979).

53. Hess, E. H. & Polt, J. M. Pupil Size in Relation to Mental Activity during Simple Problem-Solving. Science 143, 1190–1192. (2024) (Mar. 1964).

54. Naber, M. & Murphy, P. Pupillometric Investigation into the Speed-Accuracy Trade-off in a Visuo-Motor Aiming Task. Psychophysiology 57, e13499. issn: 1469-8986. (2022) (2020).

55. Richer, F. & Beatty, J. Pupillary Dilations in Movement Preparation and Execution. Psychophysiology 22, 204–207. issn: 1469-8986. (2022) (1985).

56. Van der Stigchel, S., Meeter, M. & Theeuwes, J. Eye Movement Trajectories and What They Tell Us. Neuroscience & Biobehavioral Reviews 30, 666–679. issn: 0149-7634. (2022) (Jan. 2006).

57. McPeek, R. M., Han, J. H. & Keller, E. L. Competition Between Saccade Goals in the Superior Colliculus Produces Saccade Curvature. Journal of Neurophysiology 89, 2577–2590. issn: 0022-3077. (2023) (May 2003).

58. Van der Stigchel, S. Recent Advances in the Study of Saccade Trajectory Deviations. Vision Research 50, 1619–1627. issn: 0042-6989. (2023) (Aug. 2010).

59. Spering, M. Eye Movements as a Window into Decision-Making. Annual Review of Vision Science 8, 427–448. issn: 2374-4650 (Sept. 2022).

60. Polanía, R., Krajbich, I., Grueschow, M. & Ruff, C. C. Neural Oscillations and Synchronization Differentially Support Evidence Accumulation in Perceptual and Value-Based Decision Making. Neuron 82, 709–720. issn: 0896-6273. (2024) (May 2014).

61. Mathôt, S., Siebold, A., Donk, M. & Vitu, F. Large Pupils Predict Goal-Driven Eye Movements. Journal of Experimental Psychology: General 144, 513–521. issn: 1939-2222 (2015).

62. Tkačik, G. et al. Natural Images from the Birthplace of the Human Eye. PLOS ONE 6, e20409. issn: 1932-6203. (2023) (June 2011).

63. May, J. G., Kennedy, R. S., Williams, M. C., Dunlap, W. P. & Brannan, J. R. Eye Movement Indices of Mental Workload. Acta Psychologica 75, 75–89. issn: 0001-6918. (2023) (Oct. 1990).

64. Walter, K. & Bex, P. Cognitive Load Influences Oculomotor Behavior in Natural Scenes. Scientific Reports 11, 12405. issn: 2045-2322. (2023) (June 2021).

65. Cui, M. E. & Herrmann, B. Eye Movements Decrease during Effortful Speech Listening. Journal of Neuroscience 43, 5856–5869. issn: 0270-6474, 1529-2401. (2024) (Aug. 2023).

66. Jainta, S., Vernet, M., Yang, Q. & Kapoula, Z. The Pupil Reflects Motor Preparation for Saccades - Even before the Eye Starts to Move. Frontiers in Human Neuroscience 5, 97. issn: 1662-5161. (2022) (2011).

67. Wang, C.-A. & Munoz, D. P. Differentiating Global Luminance, Arousal and Cognitive Signals on Pupil Size and Microsaccades. European Journal of Neuroscience 54, 7560–7574. issn: 1460-9568. (2022) (2021).

68. Rizzolatti, G., Riggio, L., Dascola, I. & Umiltá, C. Reorienting Attention across the Horizontal and Vertical Meridians: Evidence in Favor of a Premotor Theory of Attention. Neuropsychologia 25, 31–40. issn: 0028-3932. (2022) (Jan. 1987).

69. Deubel, H. & Schneider, W. X. Saccade Target Selection and Object Recognition: Evidence for a Common Attentional Mechanism. Vision Research 36, 1827–1837. issn: 0042-6989. (2022) (June 1996).

70. Rolfs, M. Attention in Active Vision: A Perspective on Perceptual Continuity Across Saccades. Perception 44, 900–919. issn: 0301-0066. (2022) (Aug. 2015).

71. Melcher, D. Predictive Remapping of Visual Features Precedes Saccadic Eye Movements. Nature Neuroscience 10, 903–907. issn: 1546-1726. (2023) (July 2007).

72. Bays, P. M. & Husain, M. Spatial Remapping of the Visual World across Saccades. Neuroreport 18, 1207–1213. issn: 0959-4965. (2022) (Aug. 2007).

73. Fabius, J. H., Fracasso, A., Nijboer, T. C. W. & Van der Stigchel, S. Time Course of Spatiotopic Updating across Saccades. Proceedings of the National Academy of Sciences 116, 2027–2032. (2023) (Feb. 2019).

74. Sparks, D. L. The Brainstem Control of Saccadic Eye Movements. Nature Reviews Neuroscience 3, 952–964. issn: 1471-0048. (2022) (Dec. 2002).

75. King, W. M. & Fuchs, A. F. Reticular Control of Vertical Saccadic Eye Movements by Mesencephalic Burst Neurons. Journal of Neurophysiology 42, 861– 876. issn: 0022-3077 (May 1979).

76. Curthoys, I. S., Markham, C. H. & Furuya, N. Direct Projection of Pause Neurons to Nystagmus-Related Excitatory Burst Neurons in the Cat Pontine Reticular Formation. Experimental Neurology 83, 414–422. issn: 0014-4886. (2024) (Feb. 1984).

77. Wang, C.-A., Blohm, G., Huang, J., Boehnke, S. E. & Munoz, D. P. Multisensory Integration in Orienting Behavior: Pupil Size, Microsaccades, and Saccades. Biological Psychology 129, 36–44. issn: 0301-0511. (2022) (Oct. 2017).

78. Wang, C.-A., Nguyen, K. T. & Juan, C.-H. Linking Pupil Size Modulated by Global Luminance and Motor Preparation to Saccade Behavior. Neuroscience 476, 90–101. issn: 0306-4522. (2022) (Nov. 2021).

79. Melcher, D. & Colby, C. L. Trans-Saccadic Perception. Trends in Cognitive Sciences 12, 466–473. issn: 1364-6613, 1879-307X. (2023) (Dec. 2008).

80. Van der Stigchel, S. & Hollingworth, A. Visuospatial Working Memory as a Fundamental Component of the Eye Movement System. Current Directions in Psychological Science 27, 136–143. issn: 0963-7214. (2022) (Apr. 2018).

81. Wexler, M., Mamassian, P. & Schütz, A. C. Structure of Visual Biases Revealed by Individual Differences. Vision Research 195, 108014. issn: 0042-6989. (2024) (June 2022).

82. Barbot, A., Xue, S. & Carrasco, M. Asymmetries in Visual Acuity around the Visual Field. Journal of Vision 21, 2. issn: 1534-7362. (2024) (Jan. 2021).

83. Baldwin, A. S., Meese, T. S. & Baker, D. H. The Attenuation Surface for Contrast Sensitivity Has the Form of a Witch’s Hat within the Central Visual Field. Journal of Vision 12, 23. issn: 1534-7362. (2024) (Oct. 2012).

84. Greenwood, J. A., Szinte, M., Sayim, B. & Cavanagh, P. Variations in Crowding, Saccadic Precision, and Spatial Localization Reveal the Shared Topology of Spatial Vision. Proceedings of the National Academy of Sciences 114, E3573– E3582. (2024) (Apr. 2017).

85. Chakravarthi, R., Papadaki, D. & Krajnik, J. Visual Field Asymmetries in Numerosity Processing. *Attention, Perception*, & Psychophysics 84, 2607–2622. issn: 1943-393X. (2024) (Nov. 2022).

86. Abrams, J., Nizam, A. & Carrasco, M. Isoeccentric Locations Are Not Equivalent: The Extent of the Vertical Meridian Asymmetry. Vision Research 52, 70–78. issn: 0042-6989. (2024) (Jan. 2012).

87. Ezzo, R., Winawer, J., Carrasco, M. & Rokers, B. Asymmetries in the Discrimination of Motion Direction around the Visual Field. Journal of Vision 23, 19. issn: 1534-7362. (2024) (Mar. 2023).

88. Hanning, N. M., Himmelberg, M. M. & Carrasco, M. Presaccadic Attention Depends on Eye Movement Direction and Is Related to V1 Cortical Magnification. Journal of Neuroscience 44. issn: 0270-6474, 1529-2401. (2024) (Mar. 2024).

89. Hanning, N. M., Himmelberg, M. M. & Carrasco, M. Presaccadic Attention Enhances Contrast Sensitivity, but Not at the Upper Vertical Meridian. iScience 25, 103851. issn: 2589-0042 (Feb. 2022).

90. Schütz, A. C. Interindividual Differences in Preferred Directions of Perceptual and Motor Decisions. Journal of Vision 14, 16. issn: 1534-7362. (2024) (Oct. 2014).

91. Carrasco, M., Talgar, C. P. & Cameron, E. L. Characterizing Visual Performance Fields: Effects of Transient Covert Attention, Spatial Frequency, Eccentricity, Task and Set Size. Spatial vision 15, 61. (2024) (2001).

92. Mackeben, M. Sustained Focal Attention and Peripheral Letter Recognition. Spatial vision 12, 51–72. issn: 0169-1015. (2024) (Jan. 1999).

93. Ohl, S., Kroell, L. M. & Rolfs, M. Saccadic Selection in Visual Working Memory Is Robust across the Visual Field and Linked to Saccade Metrics: Evidence from Nine Experiments and More than 100,000 Trials. Journal of Experimental Psychology: General 153, 544–563. issn: 1939-2222 (2024).

94. Greenwood, J. A., Szinte, M., Sayim, B. & Cavanagh, P. Variations in Crowding, Saccadic Precision, and Spatial Localization Reveal the Shared Topology of Spatial Vision. Proceedings of the National Academy of Sciences 114, E3573– E3582. (2024) (Apr. 2017).

95. Himmelberg, M. M., Winawer, J. & Carrasco, M. Linking Individual Differences in Human Primary Visual Cortex to Contrast Sensitivity around the Visual Field. Nature Communications 13, 3309. issn: 2041-1723. (2024) (June 2022).

96. Van Essen, D. C., Newsome, W. T. & Maunsell, J. H. R. The Visual Field Representation in Striate Cortex of the Macaque Monkey: Asymmetries, Anisotropies, and Individual Variability. Vision Research 24, 429–448. issn: 0042-6989. (2024) (Jan. 1984).

97. Benson, N. C., Kupers, E. R., Barbot, A., Carrasco, M. & Winawer, J. Cortical Magnification in Human Visual Cortex Parallels Task Performance around the Visual Field. eLife 10 (eds Meng, M., Gold, J. I., Meng, M., Aguirre, G. K. & Zuo, X.-N.) e67685. issn: 2050-084X. (2024) (Aug. 2021).

98. Himmelberg, M. M. et al. Cross-Dataset Reproducibility of Human Retinotopic Maps. NeuroImage 244, 118609. issn: 1053-8119. (2024) (Dec. 2021).

99. Silva, M. F. et al. Radial Asymmetries in Population Receptive Field Size and Cortical Magnification Factor in Early Visual Cortex. NeuroImage 167, 41–52. issn: 1053-8119. (2024) (Feb. 2018).

100. Himmelberg, M. M., Winawer, J. & Carrasco, M. Polar Angle Asymmetries in Visual Perception and Neural Architecture. Trends in Neurosciences 46, 445–458. issn: 0166-2236. (2023) (June 2023).

101. Shadmehr, R., Reppert, T. R., Summerside, E. M., Yoon, T. & Ahmed, A. A. Movement Vigor as a Reflection of Subjective Economic Utility. Trends in Neurosciences 42, 323–336. issn: 0166-2236, 1878-108X. (2024) (May 2019).

102. Voudouris, D., Schuetz, I., Schinke, T. & Fiehler, K. Pupil Dilation Scales with Movement Distance of Real but Not of Imagined Reaching Movements. Journal of Neurophysiology 130, 104–116. issn: 0022-3077. (2024) (July 2023).

103. Kool, W., McGuire, J. T., Rosen, Z. B. & Botvinick, M. M. Decision Making and the Avoidance of Cognitive Demand. Journal of Experimental Psychology: General 139, 665–682. issn: 1939-2222 (2010).

104. Kurzban, R., Duckworth, A., Kable, J. W. & Myers, J. An Opportunity Cost Model of Subjective Effort and Task Performance. Behavioral and Brain Sciences 36, 661–679. issn: 0140-525X, 1469-1825. (2024) (Dec. 2013).

105. Shenhav, A., Prater Fahey, M. & Grahek, I. Decomposing the Motivation to Exert Mental Effort. Current Directions in Psychological Science 30, 307–314. issn: 0963-7214. (2024) (Aug. 2021).

106. Shenhav, A., Botvinick, M. M. & Cohen, J. D. The Expected Value of Control: An Integrative Theory of Anterior Cingulate Cortex Function. Neuron 79, 217–240. issn: 0896-6273. (2024) (July 2013).

107. Shenhav, A. et al. Toward a Rational and Mechanistic Account of Mental Effort. Annual Review of Neuroscience 40, 99–124. issn: 0147-006X, 1545-4126. (2024) (July 2017).

108. Westbrook, A. & Braver, T. S. Cognitive Effort: A Neuroeconomic Approach. *Cognitive, Affective*, & Behavioral Neuroscience 15, 395–415. issn: 1531-135X. (2024) (June 2015).

109. Bustamante, L. A. et al. Effort Foraging Task Reveals Positive Correlation between Individual Differences in the Cost of Cognitive and Physical Effort in Humans. Proceedings of the National Academy of Sciences 120, e2221510120. (2024) (Dec. 2023).

110. Müller, T., Husain, M. & Apps, M. A. J. Preferences for Seeking Effort or Reward Information Bias the Willingness to Work. Scientific Reports 12, 19486. issn: 2045-2322. (2024) (Nov. 2022).

111. Lisi, M., Solomon, J. A. & Morgan, M. J. Gain Control of Saccadic Eye Movements Is Probabilistic. Proceedings of the National Academy of Sciences 116, 16137– 16142. (2024) (Aug. 2019).

112. Sedaghat-Nejad, E. & Shadmehr, R. The Cost of Correcting for Error during Sensorimotor Adaptation. Proceedings of the National Academy of Sciences 118, e2101717118. (2024) (Oct. 2021).

113. Petitet, P., Attaallah, B., Manohar, S. G. & Husain, M. The Computational Cost of Active Information Sampling before Decision-Making under Uncertainty. Nature Human Behaviour 5, 935–946. issn: 2397-3374. (2024) (July 2021).

114. Tatler, B. W., Brockmole, J. R. & Carpenter, R. H. S. LATEST: A Model of Saccadic Decisions in Space and Time. Psychological Review 124, 267–300. issn: 1939-1471 (2017).

115. Attneave, F. Applications of Information Theory to Psychology: A Summary of Basic Concepts, Methods, and Results vii, 120 (Henry Holt, Oxford, England, 1959).

116. Hasenstaub, A., Otte, S., Callaway, E. & Sejnowski, T. J. Metabolic Cost as a Unifying Principle Governing Neuronal Biophysics. Proceedings of the National Academy of Sciences 107, 12329–12334. (2024) (July 2010).

117. Jamadar, S., Behler, A., Deery, H. & Breakspear, P. M. The Metabolic Costs of Cognition July 2024. (2024).

118. Mathôt, S. & Vilotijević, A. Methods in Cognitive Pupillometry: Design, Preprocessing, and Statistical Analysis. Behavior Research Methods. issn: 1554-3528. (2022) (Aug. 2022).

119. Joshi, S., Li, Y., Kalwani, R. M. & Gold, J. I. Relationships between Pupil Diameter and Neuronal Activity in the Locus Coeruleus, Colliculi, and Cingulate Cortex. Neuron 89, 221–234. issn: 0896-6273. (2020) (Jan. 2016).

120. Joshi, S. & Gold, J. I. Pupil Size as a Window on Neural Substrates of Cognition. Trends in Cognitive Sciences 24, 466–480. issn: 1364-6613, 1879-307X. (2023) (June 2020).

121. Aston-Jones, G. & Cohen, J. D. An Integrative Theory Of Locus Coeruleus-Norepinephrine Function: Adaptive Gain and Optimal Performance. Annual Review of Neuroscience 28, 403–450. (2020) (2005).

122. Grujic, N., Polania, R. & Burdakov, D. Neurobehavioral Meaning of Pupil Size. Neuron 0. issn: 0896-6273. (2024) (June 2024).

123. Mazancieux, A., et al. Brainstem fMRI Signaling of Surprise across Different Types of Deviant Stimuli. Cell Reports 42. issn: 2211-1247. (2024) (Nov. 2023).

124. Schwarz, L. A. & Luo, L. Organization of the Locus Coeruleus-Norepinephrine System. Current biology: CB 25, R1051–R1056. issn: 1879-0445 (Nov. 2015).

125. Szabadi, E. Functional Neuroanatomy of the Central Noradrenergic System. Journal of Psychopharmacology 27, 659–693. issn: 0269-8811. (2020) (Aug. 2013).

126. Berridge, C. W. & Waterhouse, B. D. The Locus Coeruleus–Noradrenergic System: Modulation of Behavioral State and State-Dependent Cognitive Processes. Brain Research Reviews 42, 33–84. issn: 0165-0173. (2020) (Apr. 2003).

127. Aston-Jones, G. & Waterhouse, B. Locus Coeruleus: From Global Projection System to Adaptive Regulation of Behavior. Brain Research. 50th Anniversary Issue 1645, 75–78. issn: 0006-8993. (2020) (Aug. 2016).

128. Aston-Jones, G. & Cohen, J. D. Adaptive Gain and the Role of the Locus Coeruleus–Norepinephrine System in Optimal Performance. Journal of Comparative Neurology 493, 99–110. issn: 1096-9861. (2020) (2005).

129. Poe, G. R. et al. Locus Coeruleus: A New Look at the Blue Spot. Nature Reviews Neuroscience 21, 644–659. issn: 1471-0048. (2023) (Nov. 2020).

130. Wainstein, G., Müller, E. J., Taylor, N., Munn, B. & Shine, J. M. The Role of the Locus Coeruleus in Shaping Adaptive Cortical Melodies. Trends in Cognitive Sciences 26, 527–538. issn: 1879-307X (June 2022).

131. Dahl, M. J., Mather, M. & Werkle-Bergner, M. Noradrenergic Modulation of Rhythmic Neural Activity Shapes Selective Attention. Trends in Cognitive Sciences 26, 38–52. issn: 1364-6613. (2021) (Jan. 2022).

132. Corbetta, M., Patel, G. & Shulman, G. L. The Reorienting System of the Human Brain: From Environment to Theory of Mind. Neuron 58, 306–324. issn: 0896-6273. (2020) (May 2008).

133. Posner, M. I., Sheese, B. E., Odludaş, Y. & Tang, Y. Analyzing and Shaping Human Attentional Networks. Neural Networks. Brain and Attention 19, 1422– 1429. issn: 0893-6080. (2023) (Nov. 2006).

134. Attwell, D. & Laughlin, S. B. An Energy Budget for Signaling in the Grey Matter of the Brain. Journal of Cerebral Blood Flow & Metabolism 21, 1133–1145. issn: 0271-678X. (2024) (Oct. 2001).

135. Wiehler, A., Branzoli, F., Adanyeguh, I., Mochel, F. & Pessiglione, M. A Neuro-Metabolic Account of Why Daylong Cognitive Work Alters the Control of Economic Decisions. Current Biology 32, 3564–3575.e5. issn: 0960-9822. (2022) (Aug. 2022).

136. Castrillon, G. et al. An Energy Costly Architecture of Neuromodulators for Human Brain Evolution and Cognition. Science Advances 9, eadi7632. (2023) (Dec. 2023).

137. Gottlieb, J. Attention, Learning, and the Value of Information. Neuron 76, 281– 295. issn: 1097-4199 (Oct. 2012).

138. Kosch, T., Hassib, M., Woźniak, P. W., Buschek, D. & Alt, F. Your Eyes Tell: Leveraging Smooth Pursuit for Assessing Cognitive Workload in Proceedings of the 2018 CHI Conference on Human Factors in Computing Systems (Association for Computing Machinery, New York, NY, USA, Apr. 2018), 1–13. isbn: 978-1-4503-5620-6. (2023).

139. Siegenthaler, E. et al. Task Difficulty in Mental Arithmetic Affects Microsaccadic Rates and Magnitudes. European Journal of Neuroscience 39, 287–294. issn: 1460-9568. (2023) (2014).

140. May, J. G., Kennedy, R. S., Williams, M. C., Dunlap, W. P. & Brannan, J. R. Eye Movement Indices of Mental Workload. Acta Psychologica 75, 75–89. issn: 0001-6918. (2023) (Oct. 1990).

141. Di Stasi, L. L., Antolí, A. & Cañas, J. J. Main Sequence: An Index for Detecting Mental Workload Variation in Complex Tasks. Applied Ergonomics 42, 807– 813. issn: 0003-6870. (2024) (Nov. 2011).

142. App, E. & Debus, G. Saccadic Velocity and Activation: Development of a Diagnostic Tool for Assessing Energy Regulation. Ergonomics 41, 689–697. issn: 0014-0139 (1998).

143. Sylvestre, P. A. & Cullen, K. E. Quantitative Analysis of Abducens Neuron Discharge Dynamics during Saccadic and Slow Eye Movements. Journal of Neurophysiology 82, 2612–2632. issn: 0022-3077 (Nov. 1999).

144. Somai, R. S., Schut, M. J. & Van der Stigchel, S. Evidence for the World as an External Memory: A Trade-off between Internal and External Visual Memory Storage. Cortex; a Journal Devoted to the Study of the Nervous System and Behavior 122, 108–114. issn: 1973-8102 (Jan. 2020).

145. Van der Stigchel, S. An Embodied Account of Visual Working Memory. Visual Cognition 28, 414–419. issn: 1350-6285. (2021) (Sept. 2020).

146. Hoogerbrugge, A. J., Strauch, C., Nijboer, T. C. W. & Van der Stigchel, S. Don’t Hide the Instruction Manual: A Dynamic Trade-off between Using Internal and External Templates during Visual Search. Journal of Vision 23, 14. issn: 1534-7362. (2023) (July 2023).

147. Ballard, D. H., Hayhoe, M. M. & Pelz, J. B. Memory Representations in Natural Tasks. Journal of Cognitive Neuroscience 7, 66–80. issn: 0898-929X. (2022) (Jan. 1995).

148. Luck, S. J. & Vogel, E. K. Visual Working Memory Capacity: From Psychophysics and Neurobiology to Individual Differences. Trends in Cognitive Sciences 17, 391–400. issn: 1364-6613. (2022) (Aug. 2013).

149. Ma, W. J., Husain, M. & Bays, P. M. Changing Concepts of Working Memory. Nature Neuroscience 17, 347–356. issn: 1546-1726. (2023) (Mar. 2014).

150. Nozari, N. & Martin, R. C. Is Working Memory Domain-General or Domain-Specific? Trends in Cognitive Sciences 0. issn: 1364-6613, 1879-307X. (2024) (July 2024).

151. Paus, T. Location and Function of the Human Frontal Eye-Field: A Selective Review. Neuropsychologia 34, 475–483. issn: 0028-3932. (2024) (June 1996).

152. Bruce, C. J., Goldberg, M. E., Bushnell, M. C. & Stanton, G. B. Primate Frontal Eye Fields. II. Physiological and Anatomical Correlates of Electrically Evoked Eye Movements. Journal of Neurophysiology 54, 714–734. issn: 0022-3077. (2022) (Sept. 1985).

153. Schlag, J. & Schlag-Rey, M. Evidence for a Supplementary Eye Field. Journal of Neurophysiology 57, 179–200. issn: 0022-3077. (2024) (Jan. 1987).

154. Amiez, C. & Petrides, M. Anatomical Organization of the Eye Fields in the Human and Non-Human Primate Frontal Cortex. Progress in Neurobiology 89, 220–230. issn: 0301-0082. (2024) (Oct. 2009).

155. Sharika, K. M. et al. Proactive Control of Sequential Saccades in the Human Supplementary Eye Field. Proceedings of the National Academy of Sciences 110, E1311–E1320. (2024) (Apr. 2013).

156. Gaymard, B. et al. Effects of Anterior Cingulate Cortex Lesions on Ocular Saccades in Humans. Experimental Brain Research 120, 173–183. issn: 1432-1106. (2024) (May 1998).

157. Ruehl, R. M., Ophey, L., Ertl, M. & zu Eulenburg, P. The Cingulate Oculomotor Cortex. Cortex 138, 341–355. issn: 0010-9452. (2024) (May 2021).

158. Pouget, P. The Cortex Is in Overall Control of ‘Voluntary’ Eye Movement. Eye 29, 241–245. issn: 1476-5454. (2024) (Feb. 2015).

159. Conti, F. & Irish, M. Harnessing Visual Imagery and Oculomotor Behaviour to Understand Prospection. Trends in Cognitive Sciences 25, 272–283. issn: 1364-6613, 1879-307X. (2024) (Apr. 2021).

160. Gandhi, N. J. & Katnani, H. A. Motor Functions of the Superior Colliculus. Annual Review of Neuroscience 34, 205–231. issn: 1545-4126 (2011).

161. Sparks, D. L. Conceptual Issues Related to the Role of the Superior Colliculus in the Control of Gaze. Current Opinion in Neurobiology 9, 698–707. issn: 0959-4388. (2024) (Dec. 1999).

162. Glimcher, P. W. & Sparks, D. L. Movement Selection in Advance of Action in the Superior Colliculus. Nature 355, 542–545. issn: 1476-4687. (2024) (Feb. 1992).

163. Wurtz, R. H. & Albano, J. E. Visual-Motor Function of the Primate Superior Colliculus. Annual Review of Neuroscience 3, 189–226. issn: 0147-006X (1980).

164. Basso, M. A. & May, P. J. Circuits for Action and Cognition: A View from the Superior Colliculus. Annual Review of Vision Science 3, 197–226. issn: 2374-4650 (Sept. 2017).

165. Voogd, J. & Barmack, N. H. in Progress in Brain Research (ed Büttner-Ennever, J. A.) 231–268 (Elsevier, Jan. 2006). (2024).

166. Tanaka, M. et al. Roles of the Cerebellum in Motor Preparation and Prediction of Timing. Neuroscience. In Memoriam: Masao Ito—A Visionary Neuroscientist with a Passion for the Cerebellum 462, 220–234. issn: 0306-4522. (2024) (May 2021).

167. Takagi, M., Zee, D. S. & Tamargo, R. J. Effects of Lesions of the Oculomotor Vermis on Eye Movements in Primate: Saccades. Journal of Neurophysiology 80, 1911–1931. issn: 0022-3077. (2024) (Oct. 1998).

168. Zhang, B. et al. Transforming Absolute Value to Categorical Choice in Primate Superior Colliculus during Value-Based Decision Making. Nature Communications 12, 3410. issn: 2041-1723. (2023) (June 2021).

169. Chudasama, Y. et al. The Role of the Anterior Cingulate Cortex in Choices Based on Reward Value and Reward Contingency. Cerebral Cortex 23, 2884–2898. issn: 1047-3211. (2024) (Dec. 2013).

170. Baumann, O. et al. Consensus Paper: The Role of the Cerebellum in Perceptual Processes. The Cerebellum 14, 197–220. issn: 1473-4230. (2024) (Apr. 2015).

171. Schall, J. D. Neural Basis of Deciding, Choosing and Acting. Nature Reviews Neuroscience 2, 33–42. issn: 1471-0048. (2024) (Jan. 2001).

172. Wang, L., McAlonan, K., Goldstein, S., Gerfen, C. R. & Krauzlis, R. J. A Causal Role for Mouse Superior Colliculus in Visual Perceptual Decision-Making. Journal of Neuroscience 40, 3768–3782. issn: 0270-6474, 1529-2401. (2023) (May 2020).

173. Jun, E. J. et al. Causal Role for the Primate Superior Colliculus in the Computation of Evidence for Perceptual Decisions. Nature Neuroscience 24, 1121–1131. issn: 1546-1726. (2023) (Aug. 2021).

174. Crapse, T. B., Lau, H. & Basso, M. A. A Role for the Superior Colliculus in Decision Criteria. Neuron 97, 181–194.e6. issn: 1097-4199 (Jan. 2018).

175. Deverett, B., Koay, S. A., Oostland, M. & Wang, S. S.-H. Cerebellar Involvement in an Evidence-Accumulation Decision-Making Task. eLife 7 (eds Carey, M. R. & Ivry, R. B.) e36781. issn: 2050-084X. (2024) (Aug. 2018).

176. Basso, M. A., Bickford, M. E. & Cang, J. Unraveling Circuits of Visual Perception and Cognition through the Superior Colliculus. Neuron 109, 918–937. issn: 1097-4199 (Mar. 2021).

177. So, N. & Stuphorn, V. Supplementary Eye Field Encodes Confidence in Decisions Under Risk. Cerebral Cortex 26, 764–782. issn: 1047-3211. (2024) (Feb. 2016).

178. Yang, S.-n. & Heinen, S. Contrasting the Roles of the Supplementary and Frontal Eye Fields in Ocular Decision Making. Journal of Neurophysiology 111, 2644– 2655. issn: 0022-3077. (2024) (June 2014).

179. Padoa-Schioppa, C. & Conen, K. E. Orbitofrontal Cortex: A Neural Circuit for Economic Decisions. Neuron 96, 736–754. issn: 1097-4199 (Nov. 2017).

180. Wallis, J. D. Orbitofrontal Cortex and Its Contribution to Decision-Making. Annual Review of Neuroscience 30, 31–56. issn: 0147-006X (2007).

181. Peirce, J. et al. PsychoPy2: Experiments in Behavior Made Easy. Behavior Research Methods 51, 195–203. issn: 1554-3528. (2021) (Feb. 2019).

182. Van der Stigchel, S. & Nijboer, T. C. W. How Global Is the Global Effect? The Spatial Characteristics of Saccade Averaging. Vision Research 84, 6–15. issn: 0042-6989. (2023) (May 2013).

183. Nyström, M. & Holmqvist, K. An Adaptive Algorithm for Fixation, Saccade, and Glissade Detection in Eyetracking Data. Behavior Research Methods 42, 188– 204. issn: 1554-3528. (2022) (Feb. 2010).

184. Hayes, T. R. & Petrov, A. A. Mapping and Correcting the Influence of Gaze Position on Pupil Size Measurements. Behavior Research Methods 48, 510–527. issn: 1554-3528. (2022) (June 2016).

185. Barr, D. J. Random Effects Structure for Testing Interactions in Linear Mixed-Effects Models. Frontiers in Psychology 4. issn: 1664-1078. (2023) (2013).

186. Mathôt, S., Schreij, D. & Theeuwes, J. OpenSesame: An Open-Source, Graphical Experiment Builder for the Social Sciences. Behavior Research Methods 44, 314–324. issn: 1554-3528. (2023) (June 2012).

